# Brain Extracellular Matrix implications in multiple neurological disorders are revealed through a meta-analysis of transcriptional changes

**DOI:** 10.1101/2024.05.19.594380

**Authors:** Hagit Sadis, David Peles, Yara Hussein, Shani Stern

**Affiliations:** Sagol Department of Neurobiology, Faculty of Natural Sciences, University of Haifa, Haifa 3498838, Israel

## Abstract

Neurological disorders comprise a wide range of illnesses that may affect the central and peripheral nervous systems. Despite diverse etiologies, patients with these disorders may share symptoms.

In this study, we aimed to explore potential common mechanisms between seven neurological disorders spanning three categories: neurodegenerative diseases, neuropsychiatric disorders, and neurodevelopmental disorders, by comparing gene expression profiles and focusing on the most prominent dysregulated genes consistently reported within and across disorders. Our results demonstrate 31 genes that are commonly differentially expressed in brain cells and tissues derived from human disease models when compared to healthy controls. These genes were enriched in brain Extracellular Matrix (ECM) pathways, Growth factor binding, Response to acid chemical, and External encapsulating structure. Remarkedly, dysregulation of ECM genes was evident separately in each of the three categories of disorders. This suggests a notable distinction in the brain ECM in disease states. Furthermore, we identified that the most frequently reported genes among all disorders were *GFAP*, and *IFITM3*.

**Key Points:**

- Analysis of 41 human studies revealed 31 significantly dysregulated genes shared among seven neurological disorders when compared to healthy controls, spanning three distinct categories: Neurodegenerative diseases, Neuropsychiatric disorders, and Neurodevelopmental disorders.
- These shared Differentially Expressed Genes (DEGs) demonstrated significant enrichment for Extracellular Matrix (ECM) pathways, Growth factor binding, Response to acid chemical, Blood vessel development, and External encapsulating structure. Particularly, *SST* and *BCL6* were the most frequently reported shared DEGs.
- Notably, each of the three categories of neurological disorders exhibited significant cellular component enrichment for ECM pathways.
- In order to distinguish noise genes (false-positive genes) from disease-relevant genes, we identified the DEGs that were reported the highest number of times per disorder. *GFAP*, followed by *IFITM3*, were found to be the most reported genes.
- Furthermore, due to partially shared symptoms, we explored commonalities between Autism Spectrum Disorders (ASD) and Schizophrenia. DEGs shared between both disorders were specifically enriched with ECM pathways, External encapsulating structure, Growth factor binding, Cell adhesion molecule binding, and PI3K-Akt signaling pathway. Noteworthy, *IFITM2, HSPB1, IFITM3, HSPA1A, MKNK2, GFAP* and *COL4A1* were among the most frequently reported shared DEGs.
- The central aspects of our findings suggest a substantial distinction between the Central Nervous System (CNS) ECM in health and disease.

## Introduction

Neurological disorders are the primary cause of disability and the second most prevalent cause of death, globally. Hundreds of millions of people worldwide are affected by neurological disorders [1]. The pathological mechanisms underlying neurological disorders remain partly understood, and to this day many neurological disorders have no cure.

Neurological disorders are a heterogeneous group of illnesses that may affect both, the central and peripheral nervous systems. Despite diverse etiologies, clinical and molecular variations within and across disorders they usually share mechanistic complex involving genetic, epigenetic, environmental-induced factors, and regional brain alterations that include wide transcriptional changes that organize into vast deviations in a variety of cellular, sub-cellular, non-cellular, functional, and molecular pathways [2].

Cellular transcriptional variations in the brain of neurological patients have profound implications on cell structure and function, disturbance to homeostasis, protein accumulations and toxicity, triggering a variety of irregular pathways [2, 3]. Additionally, the intricate nature of the disease becomes apparent from the inconsistency of dysregulated genes identified from various studies for the same disorder [4]. This lack of reproducibility across studies [4] suggests that comprehensive meta-analysis can serve as a key research tool for understanding the pathophysiology of neurological disorders.

When contemplating which aspect to prioritize in researching neurological disorders, it is essential to consider that a significant portion of neurological disorders does not involve specific DNA mutations, but rather involve complex genetic variations that are not well identified. Therefore, exploring transcriptional changes rather than the DNA sequence may offer valuable insights. Gene expression can provide a unique insight into real-time events within the cell at the time of sampling the genetic material. These events may signify the response to disruptions in homeostasis, neuroinflammation, autophagy, cellular and extracellular alterations, and epigenetic modifications that all may affect gene expression. In addition to gene expression, RNA can provide data on splice variants, identify translocations, and implicate non-coding RNA molecules involved in transcriptional regulation.

Recent studies have revealed shared neuroimmune transcriptome alterations across neurological disorders, encompassing neurodegenerative diseases, neuropsychiatric disorders, and neurodevelopmental disorders [5]. Other transcriptomic studies showed dysregulation in neuron-specific modules, astrocyte- and oligodendrocyte-specific modules, and synaptic mitochondria-related modules shared among eight disorders spanning neurodegenerative diseases, and neuropsychiatric disorders [6]. A meta-study on four neurodegenerative diseases gathering genomic, transcriptomic, and proteomic data revealed 139 shared DEGs and associated pathways involving response to heat and hypoxia, positive regulation of cytokines, angiogenesis, and RNA catabolic process [7].

Our study explores three distinct groups of neurological disorders, each typical to a different stage in life: 1. Neurodegenerative diseases, including Alzheimer’s disease, Huntington’s disease, and Parkinson’s disease, characterized by progressive neuronal loss. The incidence of these diseases rises dramatically with age (usually not manifesting before the age of 30) [8]. 2. Neuropsychiatric disorders, including bipolar disorder, Major Depression, and Schizophrenia, which affect cognition, emotion, mood, and behavior; Usually, Symptoms begin in adolescence [9]. 3. Neurodevelopmental disorders, including Autism Spectrum Disorders (ASD), which are characterized by learning difficulties, attention and memory problems, and impaired social communication. Symptoms begin in the period of early development ∼ 1.5 years [10].

We conducted a comprehensive meta-analysis, comparing gene expression profiles between neurological patients and healthy controls, aiming to explore potential relationships between seven neurological disorders encompassing three categories of disorders, and targeting the most frequently reported DEGs. Such a meta-analysis, filters out noise features (false-positive genes) from disease-relevant genes within and across neurological disorders, with the overarching goal of shedding light on the underlying disease mechanisms.

## Results

### Synaptic pathways are amongst the top dysregulated pathways in Neurodegenerative diseases

When comparing gene expression of post-mortem brain tissues, neurons, and neural organoids derived from patients with neurodegenerative diseases compared to healthy controls across 16 studies, we identified 446 significant DEGs (FDR < 0.05) common to three neurodegenerative diseases: Alzheimer’s disease, Huntington’s disease, and Parkinson’s disease. We included DEGs that were reported in more than one article for each respective disease (Supplementary Table S1 presents a list of the shared DEGs).

Figure 1A displays the top 55 reported shared DEGs, sorted by the fraction of overlapping genes, in a heatmap showing the fraction of studies in which these genes were reported for each disease. The GFAP gene, coding for the Glial Fibrillary Acidic Protein, known as the major intermediate filament protein of mature astrocytes [11], had the highest ranking according to the average fraction of reporting within the context of neurodegenerative diseases (see Methods). Other prominent DEGs that were highly reported included ADARB2, AKAP12, CACNB4, CLU, GSTA4, MAF, NEFL, PLEKHG1, SCG2, GAD1, MDH1, SNAP25, and STMN2. The shared DEGs among the diseases are depicted in a Venn Diagram (Fig. 1B).

**Figure 1.**
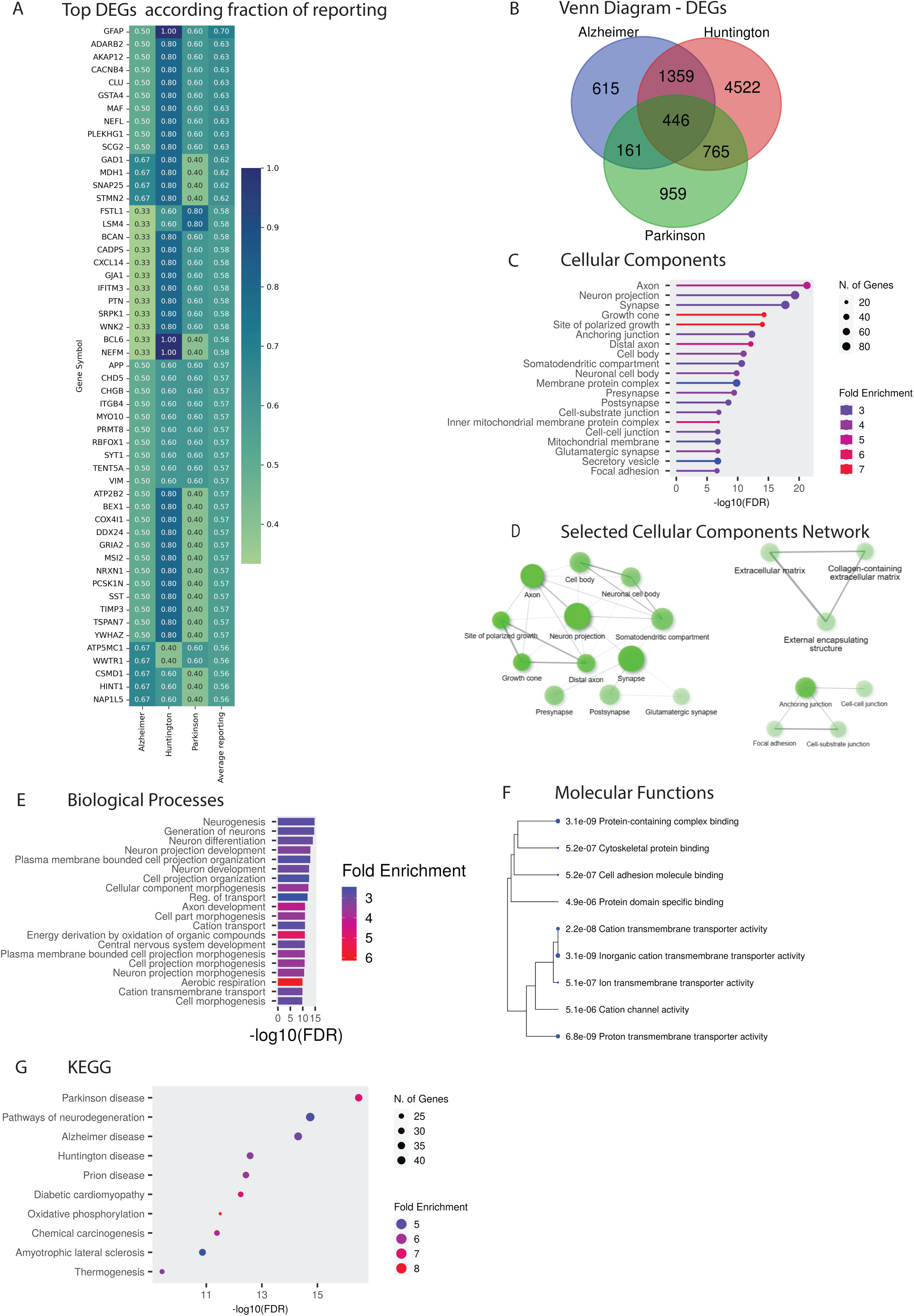
Synaptic pathways are amongst the top dysregulated pathways in Neurodegenerative diseases. **A**. A Venn Diagram showing the DEGs in the three neurodegenerative diseases: Alzheimer’s disease, Huntington’s disease, and Parkinson’s disease, compared to healthy controls that appeared in more than one study for each disease. **B.** A heatmap illustrating the top 55 common significantly reported DEGs (FDR<0.05) sorted by the average fraction of overlapping genes, depicting the fraction of studies in which these genes were reported for each disease compared to healthy controls. **C**. The top 20 significant cellular components (FDR<0.05) presenting enrichment for the shared DEGs between the three neurodegenerative diseases compared to healthy controls, which were identified in more than one study for each disease. **D**. Selected cellular component networks, depicting the interactions among significant cellular components (FDR<0.05) for shared DEGs across three neurodegenerative diseases compared to healthy controls. These DEGs were identified in more than one study for each disease. **E**. The top 20 significant biological processes (FDR<0.05) presenting enrichment for the shared DEGs between the three neurodegenerative diseases compared to healthy controls, which were identified in more than one study for each disease. **F**. The top 9 significant molecular function pathways enrichment tree (FDR<0.05) for the shared DEGs between the three neurodegenerative diseases compared to healthy controls, which were identified in more than one study for each disease. **G**. The top 10 significant KEGG pathways (FDR<0.05) presenting enrichment for the shared DEGs between the three neurodegenerative diseases compared to healthy controls, which were identified in more than one study for each disease.

The analysis of the cellular components enrichment for the shared DEGs between neurodegenerative diseases revealed several pathways including Axon, Neuron projection, Synapse, Somatodendritic compartment, Neuronal cell body, Presynapse, Postsynapse, Glutamatergic synapse, Anchoring junction, Extracellular matrix, Collagen-containing extracellular matrix, and more (Fig. 1C, D - cellular components, Supplementary Table S2 presents a complete list of the cellular component pathways). When analyzing the enrichment for biological processes, we found several pathways including Neurogenesis, Generation of neurons, Neuron differentiation, and Neuron projection development (Fig. 1E – biological processes). The most significant molecular functions enrichment included Protein-containing complex binding, Inorganic cation transmembrane transporter activity, and Proton transmembrane transporter activity (Fig. 1F - molecular functions). KEGG pathways were mostly enriched for Pathways of neurodegeneration, Parkinson’s disease, Alzheimer’s disease, Huntington’s disease, Prion diseases, Diabetic cardiomyopathy, and Amyotrophic lateral sclerosis (Fig. 1G - KEGG).

Furthermore, we conducted individual analyses for each of the three neurodegenerative diseases, described next briefly (see Supplementary S3, and extended figures 1-3 for detailed analysis).

### Alzheimer’s disease involves dysregulation of synaptic pathways, accompanied by abnormalities in the AHNAK and SCG5 genes

Alzheimer’s disease’s top DEGs, sorted by the percentage of overlapping genes among studies, were AHNAK and SCG5 (recurring in 83.3% of relevant studies) (Extended Figures 1A). Extended Figures 1B, C, D, and E display the GO analysis for 2,581 significant DEGs (FDR < 0.05) which were identified in more than one Alzheimer’s disease study out of the six relevant studies, all are coding genes (Extended Fig. 1F). These DEGs demonstrated significant enrichment mostly for Synaptic pathways including Glutamatergic synapse, Neurogenesis, Cytoskeletal protein binding, Cell adhesion molecule binding, and Pathways of neurodegeneration (see Supplementary S3 for the detailed analysis).

SCG5, coding for secretogranin V, also known as 7B2, is a secreted chaperone protein located in the extracellular region and nucleus, that prevents the aggregation of other secreted proteins, including proteins that are associated with neurodegenerative and metabolic disease. Chaplot et al. suggested that the potent chaperone action of SCG5 and its presence in a variety of extracellular protein deposits could have significance in neurodegenerative proteostatic mechanisms [12] [13] [14]. AHNAK is a large (700 kDa) structural scaffold protein, it has an essential role in developmental myelination processes, neuronal plasticity, and events related to neurodegeneration [13]. AHNAK was proposed as a potential biomarker of aging-related neurodegeneration [14].

### Parkinson’s disease involves dysregulation of mitochondrial-related pathways

Parkinson’s disease’s top DEGs, sorted by the percentage of overlapping genes among studies, were ACADVL, COL1A1, COL1A2, COL3A1, CPNE1, EEA1, FSTL1, GLIPR1, IFI16, KIAA1755, LSM4, NCAPD2, PAPPA, POU3F3, PTGDS, and TMEM107 (recurring in 80% of relevant studies) (Extended Figures 2A). Extended Figures 2B, C, D, and E display the GO analysis for 2,333 significant DEGs (FDR < 0.05) which were identified in more than one Parkinson’s disease study out of the five relevant studies, almost all are coding genes, and very few are of type long non-coding RNA (lncRNA) (Extended Fig. 2F). These DEGs demonstrated significant enrichment primarily in Mitocondrial-related pathways, Synaptic pathways, Neurogenesis, Oxidative phosphorylation, and RNA binding (see Supplementary S3 for detailed analysis). Notably, mitochondrial dysfunction was previously recognized as a significant contributor to the development of both monogenic and idiopathic Parkinson’s disease [15].

### Huntington’s disease – dysregulation of urea metabolism

In Huntington’s disease, the top DEGs, sorted by the percentage of overlapping genes among studies, were BCL6, CEBPD, CRYM, FKBP5, GFAP, HTR2C, NEFM, PLOD2, and SLC14A1 (recurring in all five relevant studies) (Extended Figures 3A). Protein-protein interaction (PPIs) network enrichment shows interactions between all nine top reported DEGs except the SLC14A1 gene (p <0.0023) (Extended Fig. 3B – Top genes PPIs networks). Surprisingly, SLC14A1, coding to solute carrier family 14 member 1 (Kidd blood group), is responsible for mediating the transport of urea. It has been reported that abnormal urea metabolism might serve as the initial biochemical disruption triggering neuropathogenesis in Huntington’s disease [16]. Extended Figures 3C, D, E, F, and G display the GO analysis for 2,898 significant DEGs (FDR < 0.05), which were identified in more than two Huntington’s disease studies out of the five relevant studies, all are coding genes (Extended Fig. 3C). These DEGs demonstrated significant enrichment primarily in Synaptic pathways including Glutamatergic synapse, Neurogenesis, Cytoskeletal protein binding, and Circadian entrainment (see Supplementary S3 for detailed analysis).

Significantly, Bailus et al. have conclusively shown that FKBP5 is involved in the progression of Huntington’s disease. Inhibiting FKBP5 in human-derived neural stem cells from individuals with Huntington’s disease, as well as in diverse mouse models of HD, resulted in a decrease in mutant Huntingtin (HTT) gene expression levels. Their findings propose a model wherein the reduction of FKBP5 levels or activity results in an increased autophagy process. This discovery positions FKBP5 as a potential therapeutic target [17].

### Prominent Dysregulation of ECM pathways in Neuropsychiatric disorders

When comparing gene expression of post-mortem brain tissues and neurons derived from neuropsychiatric patients and healthy controls across 18 studies, we identified 107 significant DEGs (FDR < 0.05) common to three neuropsychiatric disorders: Schizophrenia, Major Depression, and bipolar disorder (Supplementary Table S1 - a list of the shared DEGs).

Figure 2A displays the top 30 reported shared DEGs, sorted by the fraction of overlapping genes, in a heatmap showing the fraction of studies in which these genes were reported. The most prominent genes, according to the average fraction of reporting within the context of neuropsychiatric disorders, were HLA-A, TM4SF1, and IFI6 followed by COL4A1, SST, NR2F2, TNFRSF11B, RGS1, and FAM118A gene. The shared DEGs among the disorders are depicted in a Venn Diagram (Figure 2B). Analysis of the cellular component enrichment of DEGs shared between neuropsychiatric disorders revealed the following dysregulated pathways: Extracellular Matrix, External encapsulating structure, Collagen-containing extracellular matrix, and Endoplasmic reticulum lumen (Fig. 2C - cellular components, Supplementary Table S2 –a list of the cellular components pathways). When analyzing the enrichment for biological processes we found that the most significant pathways were Tube development, Blood vessel development, Anatomical structure formation involved in morphogenesis, Circulatory system development, Vasculature development, Tube morphogenesis, and Blood vessel morphogenesis (Fig. 2D – biological processes). Molecular Functions were enriched for Growth factor binding, Signaling receptor binding, and Signaling receptor regulator activity (Fig. 2E - molecular functions).

**Figure 2.**
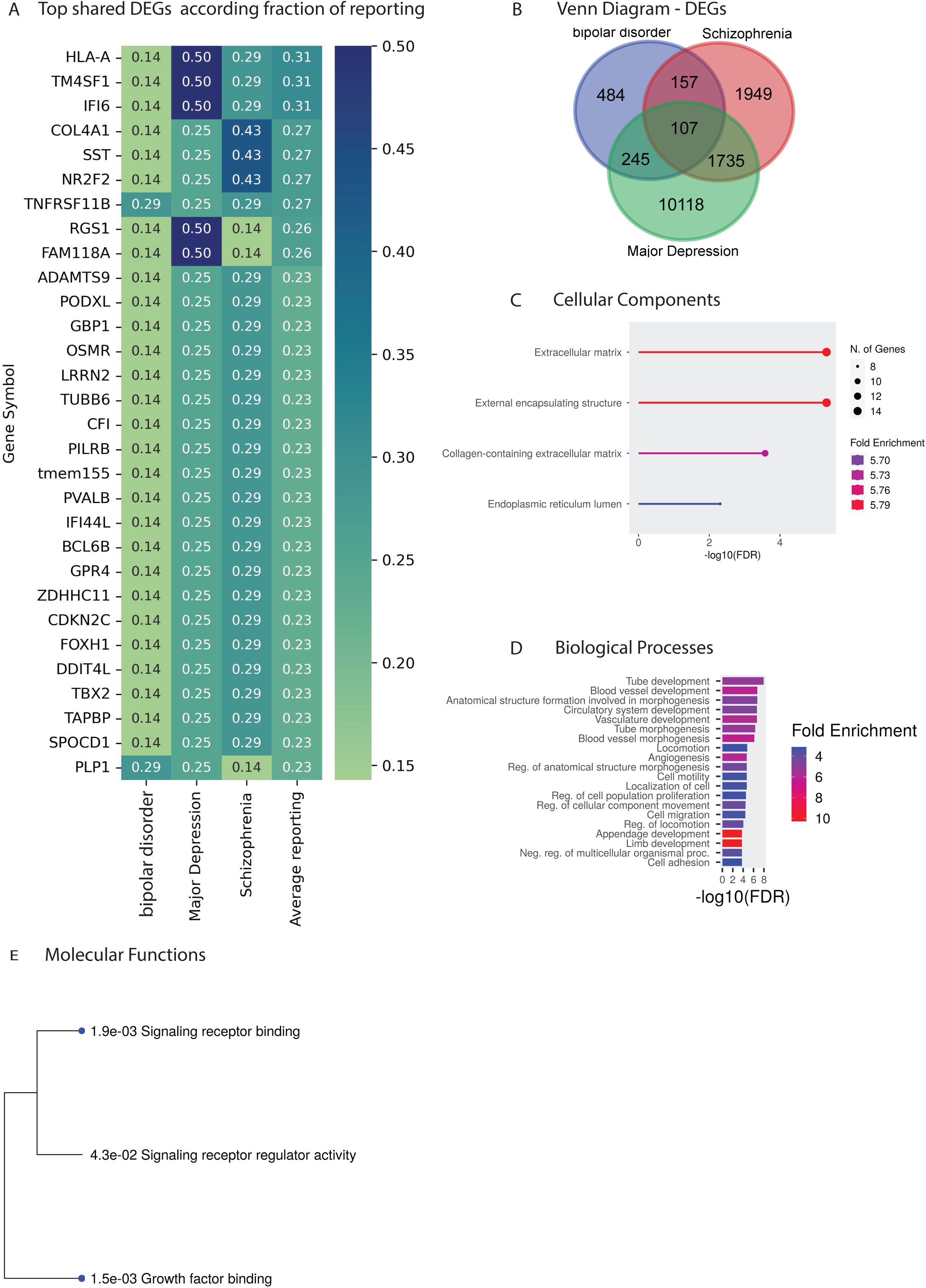
Prominent Dysregulation of ECM pathways in Neuropsychiatric disorders. **A**. A Venn Diagram representing the identified DEGs of the three neuropsychiatric disorders: bipolar disorder, Major Depression, and schizophrenia, compared to healthy controls. **B**. A heatmap illustrating the top 30 common significant reported DEGs (FDR<0.05) sorted by the average fraction of overlapping genes, depicting the fraction of studies in which these genes were reported for each disorder compared to healthy controls. **C**. The significant cellular components (FDR<0.05) presenting enrichment for the shared DEGs between the three neuropsychiatric disorders compared to healthy controls. **D**. The top 20 significant biological processes (FDR<0.05) presenting enrichment for the shared DEGs between the three neuropsychiatric disorders compared to healthy controls. **E**. The significant molecular function pathways enrichment tree (FDR<0.05) for the shared DEGs between the three neuropsychiatric disorders compared to healthy controls.

Furthermore, we conducted individual analyses for each of the three neuropsychiatric disorders, described briefly below (see Supplementary S4, and extended figures 4-6 for detailed analysis).

### Schizophrenia

Schizophrenia’s top DEGs, sorted by the percentage of overlapping genes, were HSPA1A, IFITM2, and IFITM3 (recurring in 71.4% of relevant studies) followed by IFITM1, HSPB1, HSPA1B, MKNK2, MEG3, JMJD6, (recurring in 57.1% of relevant studies) (see Extended Figures 4A). Extended Figures 4B, C, D, and E display the GO analysis for 3,955 significant DEGs (FDR < 0.05) which were identified from seven Schizophrenia studies. These DEGs demonstrated significant enrichment primarily in Synaptic pathways, Circulatory system development, Neurogenesis, Cell adhesion molecule binding, and Signaling receptor binding (see Supplementary S4 for detailed analysis). Remarkably, Stern et al. [18] showed that hippocampal neurons derived from monozygotic twins discordant for schizophrenia, differ in synapse-related genes and synaptic activity.

### Major Depression

In Major Depression, the FOS gene had the highest ranking (recurring in 75% of the relevant studies) when sorted by the percentage of overlapping genes obtained from four studies, followed by an additional 42 DEGs (recurring in 50% of the relevant studies) (see Extended Figures 5A). We identified 12,221 significant DEGs (FDR < 0.05) from four Major Depression studies. Given the substantial number of dysregulated genes, Extended Figures 5B, C, D, and E display the GO analysis for the 43 significant DEGs (FDR < 0.05) which were identified in more than one Major Depression study. These DEGs demonstrated significant enrichment primarily in Post-regulation of cell death, Skeletal muscle cell differentiation, DNA-binding transcription activator activity, and IL-17 signaling pathway (see Supplementary S4 for detailed analysis). The IL-17 (interleukin 17) family, comprising six members (IL-17A-F), is a subset of cytokines crucially involved in autoimmune diseases. Notably, high levels of IL-17 have been detected in individuals with depression [19].

### Bipolar disorder demonstrates substantial variability among patients

In bipolar disorder we identified 13 significant DEGs (FDR<0.05) that appeared in more than one bipolar disorder study out of the seven relevant studies examined. These genes that exhibit recurrence included CA12, ETNPPL, GNG12, HTR7, IL4R, MLC1, PLP1, RASGRP1, SENP6, SLC35F1, TGIF1, TNFRSF11B, and TRAF3IP2 (see Extended Figures 6A), a significantly lower overlapping gene list compared to the other disorders. One possible explanation for the low recurrence could be the substantial variability observed among bipolar patients, encompassing distinct cellular parameters between lithium-responsive patients and non-responsive patients, leading to the emergence of discrete neuronal subgroups [20, 21]]23,22[. Additionally, another study showed that distinct epigenetic mechanisms influencing gene expression categorize individuals with bipolar disorder into two groups: those with suicidal tendencies and those with non-suicidal tendencies [24]. Go analysis reveals that the 993 significant DEGs (FDR < 0.05) identified from seven bipolar disorder studies showed significant enrichment primarily in ECM pathways, Animal organ morphogenesis, Neuroactive ligand-receptor interaction, and PI3K-Akt signaling pathway (see Extended Figures 6B, C, D, E, F and Supplementary S4 for detailed analysis).

### Prominent Dysregulation of brain ECM in Neurodevelopmental disorders

When comparing gene expression of post-mortem brain tissues, neurons, and neural organoids derived from neurodevelopmental patients and healthy controls across seven studies, we identified 1,028 significant DEGs (FDR < 0.05). These DEGs were reported in more than one article for each respective disorder (Supplementary Table S1 presents a list of the shared DEGs).

Figure 3A displays the 42 top reported DEGs, sorted by the fraction of overlapping genes, in a heatmap showing the fraction of studies in which these genes were reported. The most prominent genes that were highly reported were: GFAP, ADM, CEBPD, ITPKB, MSN, and S100A10, each recurring in five out of the seven relevant studies. Analysis of the cellular component enrichment reveals that DEGs shared between neurodevelopmental disorders are significantly enriched with Extracellular Matrix, External encapsulating structure, Collagen-containing extracellular matrix, Synapse, Neuron Projection, and more (Fig. 3B - cellular components). When analyzing the enrichment for biological processes, we found that the most significant pathways were: Extracellular matrix organization, Extracellular structure organization, and External encapsulating structure organization (Fig. 3C – biological processes). Molecular Functions enrichment included Extracellular matrix structural constituent, Extracellular matrix binding, Glycosaminoglycan binding, signaling receptor binding, and Growth factor binding, among others (Fig 3D – molecular functions). KEGG pathways were enriched mostly for MAPK signaling pathways, and PI3K-Akt signaling pathway. (Fig 3E – KEGG).

**Figure 3.**
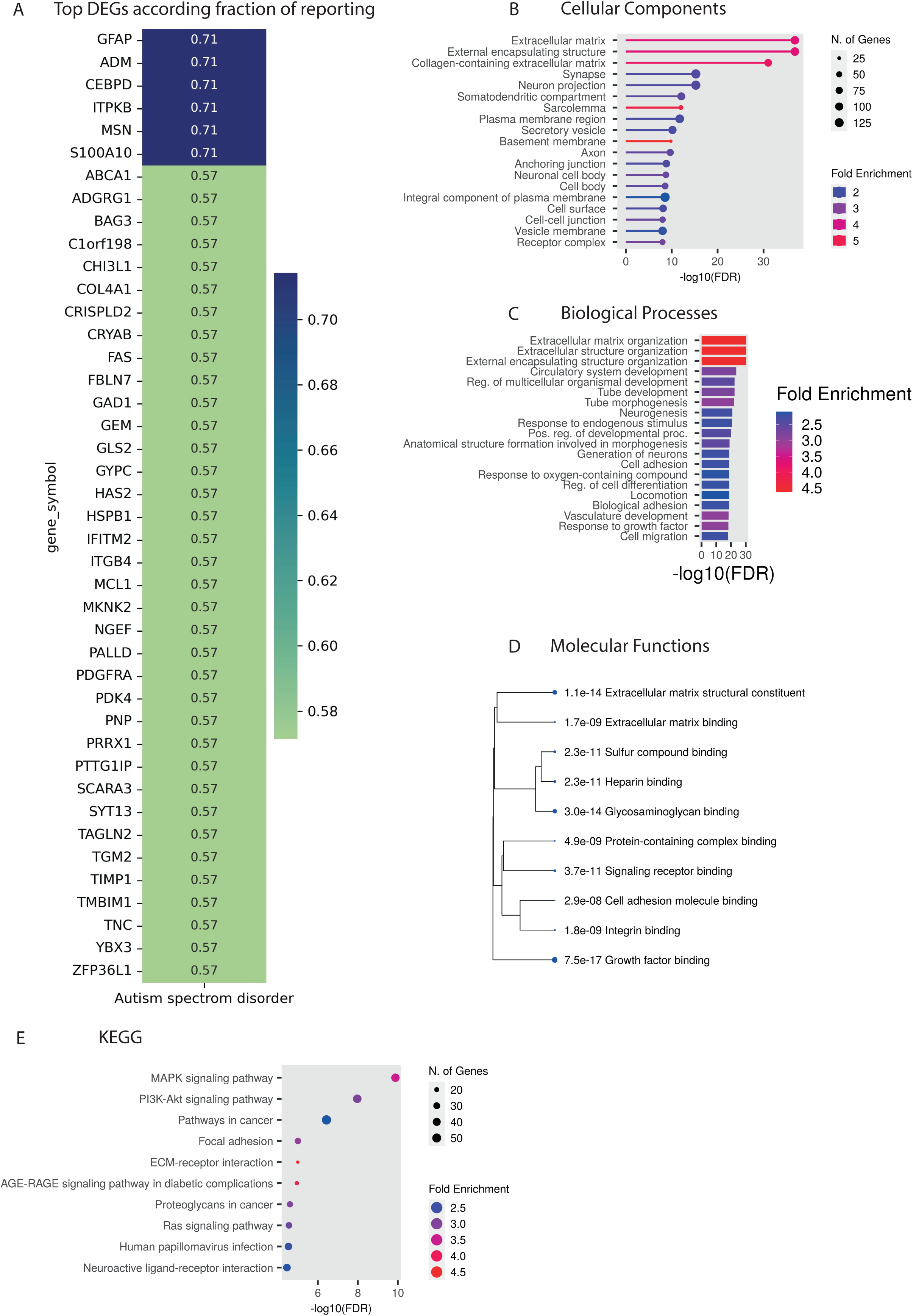
Prominent Dysregulation of brain ECM in Neurodevelopmental disorders. A. A heatmap illustrating the top 42 common significant reported DEGs (FDR<0.05) of neurodevelopmental disorders, sorted by the fraction of overlapping genes, depicting the fraction of studies in which these genes were reported compared to healthy controls. B. The top 20 significant cellular components (FDR<0.05), presenting enrichment for the shared DEGs between the neurodevelopmental disorders compared to healthy controls, which were identified in more than one study. C. The top 20 significant biological processes (FDR<0.05) presenting enrichment for the shared DEGs between neurodevelopmental disorders compared to healthy controls, which were identified in more than one study. G. The top 10 significant molecular function pathways enrichment tree (FDR<0.05) for the shared DEGs between the neurodevelopmental disorders compared to healthy controls, which were identified in more than one study. H. The top 10 significant KEGG pathways (FDR<0.05) presenting enrichment for the shared DEGs between the neurodevelopmental disorders compared to healthy controls, which were identified in more than one study.

### ECM pathways are highly dysregulated in Neurological disorders

When seeking common dysregulated genes across all three categories of disorders, which encompass neuropsychiatric disorders (bipolar disorder, Major Depression, and Schizophrenia), neurodegenerative diseases (Alzheimer’s disease, Huntington’s disease, and Parkinson’s disease), and neurodevelopmental disorders (ASD) across 41 studies, we identified 31 significantly shared DEGs (FDR<0.05). Figure 4A displays 31 common DEGs, sorted by the fraction of overlapping genes, in a heatmap showing the fraction of studies in which these genes were reported. The most prominent genes that were highly reported were: SST, BCL6, PRMT8, NFIX, COL1A2, SAT1, MT1E, SPON1, PODXL, CCN2, and HLA-E. Significantly, SST was identified in 80% of Huntington’s disease studies and 50% of Alzheimer’s disease studies analyzed. Figure 4B illustrates DEGs intersections among the three categories of disorders and Figure 4C lists some of the overlapping genes between the different groups.

**Figure 4.**
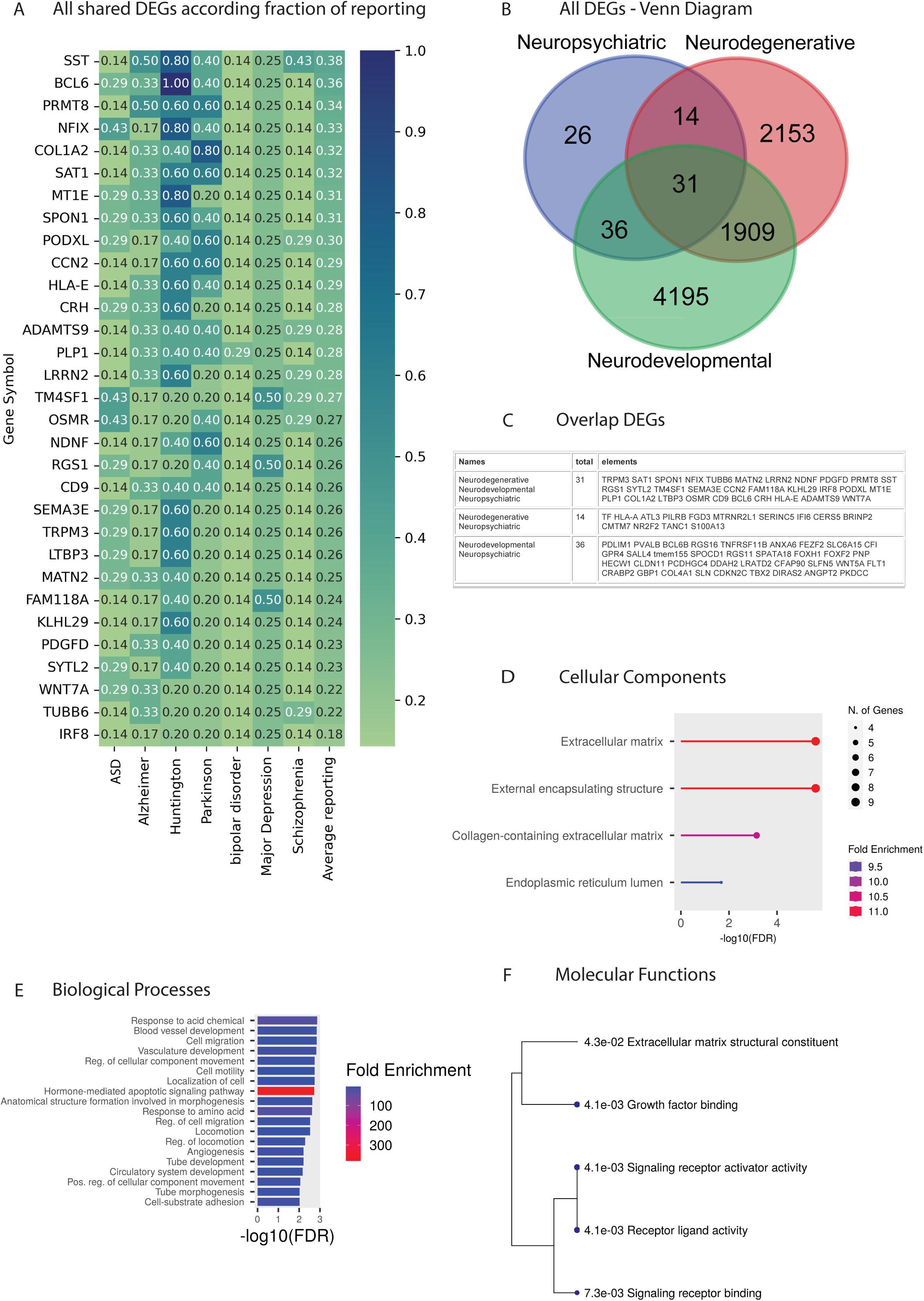
ECM pathways are highly dysregulated in Neurological disorders. **A**. A heatmap illustrating the 31 common significant reported DEGs (FDR<0.05) sorted by the average fraction of overlapping genes, among 7 neurological disorders spanning three categories of disorders, depicting the fraction of studies in which these genes were reported for each disorder compared to healthy controls. **B**. A Venn Diagram representing the identified DEGs of the three categories of neurological disorders: neurodegenerative diseases, neuropsychiatric disorders, and neurodevelopmental disorders, compared to healthy controls. **C**. The significant cellular components (FDR<0.05) presenting enrichment for the shared DEGs between the three categories of disorders compared to healthy controls. **D**. The top 20 significant biological processes (FDR<0.05) presenting enrichment for the shared DEGs between the three categories of disorders compared to healthy controls. **E**. The significant molecular function pathways enrichment tree (FDR<0.05) for the shared DEGs between the three categories of disorders compared to healthy controls.

Analysis of the cellular component enrichment reveals that DEGs shared among the three categories of neurological disorders were enriched for the following pathways: Extracellular matrix, External encapsulating structure, Collagen-containing extracellular matrix, and Endoplasmic reticulum lumen (Fig. 4D – cellular components). When analyzing the enrichment for the biological processes we found that among the most significant pathways were Response to acid chemical, Blood vessel development, Cell migration, Vasculature development, and Hormone mediated apoptotic signaling pathways (Fig. 4E – biological processes). Molecular Functions enrichment included: Extracellular matrix structural constituent, Growth factor binding, Signaling receptor activator activity, Receptor ligand activity, and Signaling receptor binding (Fig. 4F – molecular Functions).

### GFAP and IFITM3 gene’s role in neurological disorders

We next sought the most dysregulated genes within each of the disorders by the means of repetition between different reported studies.The GFAP gene followed by IFITM3, showed the highest ranking according to the average fraction of reporting, for the seven disorders (Fig. 5A – top reported DEGs, Supplementary Table S5 – DEGs Summary). Other highly reported DEGs were: HSPA1A, MKNK2, ITGB4, SST, IFITM2, GAD1, GSTA4, SCG2, CEBPD, BCL6, MDH1, FOS, CLU, ZEP36L1, and HSPB1.

**Figure 5.**
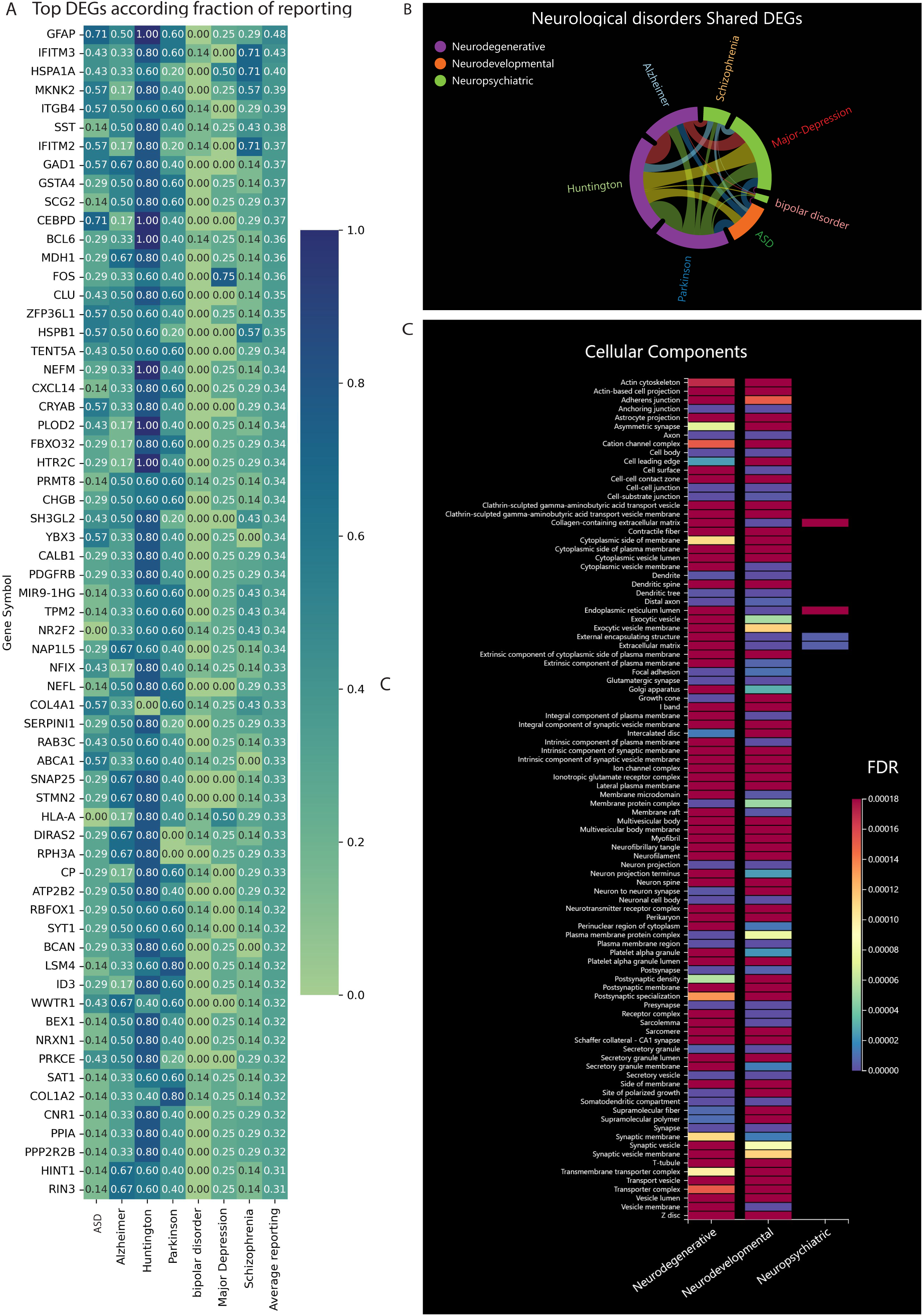
GFAP and IFITM3 gene’s role in Neurological disorders. **A**. A heatmap illustrating the most significant reported DEGs (FDR<0.05) sorted by the average fraction of overlapping genes, among seven neurological disorders (not necessarily shared among all) spanning three groups of disorders, depicting the fraction of studies in which these genes were reported for each disorder compared to healthy controls. **B**. Shared genes between seven neurological disorders, where the length of the outer arch indicates the number of DEGs identified for a disorder and the thickness of the connecting lines reflects the number of shared DEGs between each pair of disorders. **C**. Cellular component enrichment for the 3 groups of neurological disorders. Notably, only 4 cellular components were common across all 3 groups, including the Extracellular matrix, External encapsulating structure, Collagen-containing extracellular matrix, and Endoplasmic reticulum lumen.

Figure 5B presents the number of DEGs identified for each disorder and the number of shared dysregulated genes between each pair of disorders. Significantly, Huntington’s disease and Parkinson’s disease exhibit the highest overlap with 6,714 shared DEGs, followed by Huntington’s disease and Alzheimer’s disease with 6,644 shared DEGs. Among two disorders originating from distinct neurological groups, Major Depression and Huntington’s disease exhibited the most significant overlap with 6,245 shared DEGs, while bipolar disorder and Schizophrenia exhibit the lowest overlap with 264 shared DEGs (see Supplementary Table S1 – a list of the shared DEGs).

Figure 5C displays the cellular component enrichment for the three groups of neurological disorders. Notably, only four cellular components were common across all three groups, including the Extracellular matrix, External encapsulating structure, Collagen-containing extracellular matrix, and Endoplasmic reticulum lumen. Certain biological process pathways were common among multiple neurological disorder groups, including Anatomical structure formation involved in morphogenesis, Cell adhesion, Cell migration, Circulatory system development, Generation of Neurons, Locomotion, Neurogenesis, Tube development, Tube morphogenesis, Vasculature development, and more (Extended Fig. 7A – biological processes).

### ECM gene dysregulation is common between ASD and Schizophrenia

The negative symptoms of Schizophrenia resemble many ASD symptoms such as, impaired social communication, social cognition deficits, and reduced emotional expression [25]. Both ASD and Schizophrenia arise as a result of strong genetic and environmental risk factors that interact in complex ways. [26]. In addition, the neurodevelopmental hypothesis of Schizophrenia implicates abnormalities or disruptions in neural growth during embryonic development, subtle cytoarchitectural changes in the brain, and changes in the gross morphology of the brain, such as enlarged ventricles, widened sulci, and loss of normal asymmetry [27]. A Genome-wide association study (GWAS) analysis revealed that 74% of the genes most strongly linked to ASD were also associated with Schizophrenia [25]. Furthermore, Intellectual disabilty and Schizophrenia co-occur much more frequently than would be expected by chance [26, 28].

When looking for transcriptional commonalities between ASD and Schizophrenia, we identified 145 significant shared DEGs (FDR<0.05) that were reported in more than one study for each disorder (Supplementary Table S1 – a list of the shared DEGs). Figure 6A illustrates the DEGs intersections among the two disorders in a Venn diagram, and Figure 6B displays the 21 top reported shared DEGs in a heatmap sorted by the fraction of studies in which these genes were reported. The gene IFITM2 showed the highest ranking according to the average fraction of reporting. IFITM2, encoding for the interferon-induced transmembrane protein 2, is involved in immune response. Other prominent DEGs were HSPB1, IFITM3, HSPA1A, MKNK2, GFAP, COL4A1, TGM2, BAG3, and CEBPD. Analysis of the cellular component enrichment revealed that the most significant components included the Extracellular matrix, External encapsulating structure, Collagen-containing extracellular matrix, Synapse, and Glutamatergic synapse (Fig. 6C - cellular components). When analyzing the enrichment for the biological processes, the most significant pathways included Extracellular matrix organization, Extracellular structure organization, External encapsulating structure organization, and Response to endogenous stimulus (Fig. 6D – biological processes). Molecular Functions enrichment’s most significant functions included Growth factor binding, Extracellular matrix binding, Cell adhesion molecule binding, Integrin binding, and more (Fig. 6E - molecular functions). KEGG pathways were mostly enriched for PI3K-Akt signaling pathway, Human papillomavirus infection, MAPK signaling pathway, Focal adhesion, and ECM-receptor interaction (Fig. 6F – KEGG).

**Figure 6.**
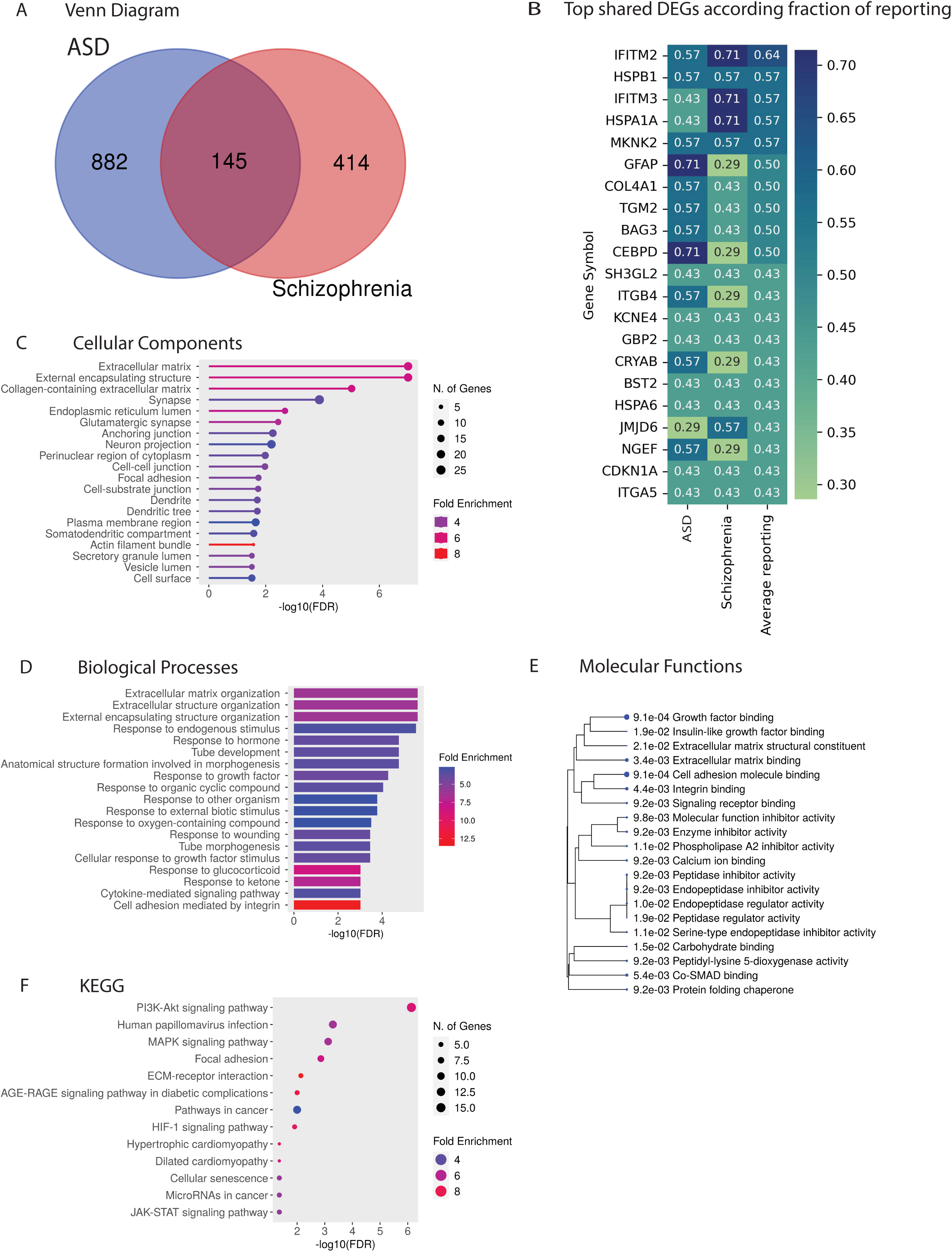
ECM gene dysregulation is common between ASD and Schizophrenia. **A**. A Venn Diagram representing the identified DEGs of the two disorders: Autism spectrum disorder (ASD) and Schizophrenia that appeared in more than one study for each disorder. **B.** A heatmap illustrating the top 21 common significant reported DEGs (FDR<0.05) sorted by the average fraction of overlapping genes, depicting the fraction of studies in which these genes were reported for each disorder, compared to healthy controls. **C**. The top 20 significant cellular components (FDR<0.05) presenting enrichment for the shared DEGs between the two disorders compared to healthy controls, which were identified in more than one study for each disorder. **D**. The top 20 significant biological processes (FDR<0.05) presenting enrichment for the shared DEGs between the two disorders compared to healthy controls, which were identified in more than one study for each disorder. **E**. The top 20 significant molecular function pathways enrichment tree (FDR<0.05) for the shared DEGs between the two disorders compared to healthy controls, which were identified in more than one study for each disorder. **F**. The top 13 significant KEGG pathways (FDR<0.05) presenting enrichment for the shared DEGs between the two disorders compared to healthy controls, which were identified in more than one study for each disorder.

## Methods

We analyzed 3 groups of neurological disorders, which encompass Neuropsychiatric disorders (bipolar disorder, Major Depression, and Schizophrenia), Neurodegenerative diseases (Alzheimer’s disease, Huntington’s disease, and Parkinson’s disease), and Neurodevelopmental disorders (Autism spectrum disorders) from a total of 41 human studies including 828 patients and 628 healthy controls. For each disorder, we compared gene expression of post-mortem brain tissues, neurons and neural organoids derived from patients and healthy controls and identified significant DEGs with a false discovery rate (FDR) < 0.05. Figure 7A illustrates the classification of tested models obtained from the studies analyzed. Figure 7B presents the categorization of studies based on the tested model and the number of studies used for each category of neurological disorders, while Figure 7C displays the classification of the tested models and the number of studies used for each disorder separately.

**Figure 7.**
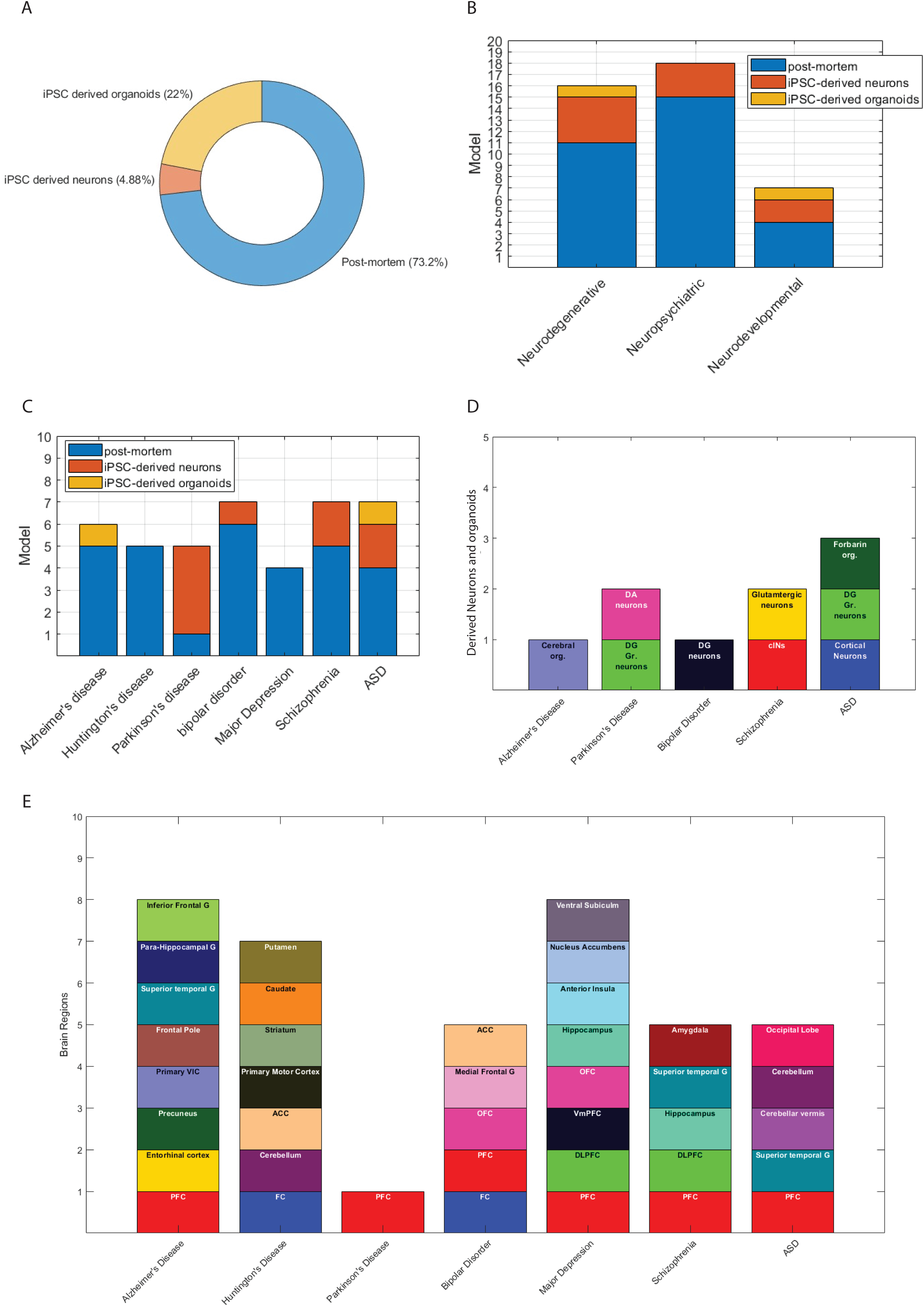
Classification of model types. **A**. The data from 41 articles encompassing seven neurological disorders were classified into three model categories: Post-mortem tissues (n=30), iPSC-derived neurons (n=9), and iPSC-derived organoids (n=2). **B**. The graphical representation depicts the categorization of studies based on the tested model for each group of neurological disorders. Three model types were considered: Post-mortem tissues, iPSC-derived neurons, and iPSC-derived organoids. The bars correspond to the number of studies conducted for each category of disorders: neurodegenerative diseases (n=16), neuropsychiatric disorders (n=18), and neurodevelopmental disorders (n=7). **C**. The studies were categorized based on the tested model for each neurological disorder. Each bar’s height corresponds to the number of studies conducted for each disorder: Alzheimer’s disease (n=6), Huntington’s disease (n=5), Parkinson’s disease (n=5), bipolar disorder (n=7), Major Depression (n=4), Schizophrenia (n=7), and autism spectrum disorders (n=7). **D**. Categorization of iPSC-derived neurons and neuronal organoids for each disorder. Abbreviations: DA-dopaminergic, DG-dentate gyrus. **E**. Classification of post-mortem tissue samples for each disorder. Abbreviations: G - Gyrus, VIC - visual cortex, FC - frontal cortex, PFC - prefrontal cortex, ACC - anterior cingulate cortex, OFC- orbitofrontal cortex, vmPC - ventromedial prefrontal cortex, DLPFC - dorsolateral prefrontal cortex

### Literature search

We used Google Scholar and the PubMed Central (PMC) archive of biomedical and life sciences journal literature (https://www.ncbi.nlm.nih.gov/pmc/) and searched for: [the name of the disorder], ’human’, ’gene expression’, or ’DEGs’ or ’RNA-Sequence’, which resulted in 80 studies. We next excluded studies that used peripheral blood, Neural Progenitor cells (NPCs), Lymphoblastoid cell lines (LCLs), animal models, and studies that demonstrated DEGs that didn’t exhibit significant FDR. Ultimately, we retained 41 studies [29–69]. Table 1 displays the list of studies used in this meta-analysis and contains details on the model examined, methods used for identifying DEGs, the number of DEGs identified, and the number of patients and healthy controls who participated in the study.

### The data used for the analysis

Induced Pluripotent Stem Cells (iPSCs) were differentiated into dentate gyrus neurons, cortical neurons, cortical interneurons (cINs), dopaminergic (DA) neurons, and glutamatergic neurons. iPSC-derived organoids included 3-D cerebral organoid models, and forebrain organoids containing a range of embryonic prefrontal cortical cell types as shown in Figure 7D.

Post-mortem tissues were obtained from diverse brain regions, including the frontal cortex, prefrontal cortex (PFC), dorsal lateral PFC (DLPFC), ventromedial PFC, orbitofrontal cortex (OFC), entorhinal cortex, precuneus (PREC), primary visual cortex (VIC), frontal pole (FP), hippocampus, superior temporal gyrus (STG), para-hippocampal gyrus (PHG), inferior frontal gyrus (IFG), medial frontal gyrus, cerebellar vermis, cerebellar, occipital lobe, anterior cingulate cortex (ACC), primary motor cortex, striatum, caudate, putamen, anterior insula, nucleus accumbens, ventral subiculum, and amygdala as shown in Figure 7E.

### Analysis of gene expression

The methods employed in the analyzed studies to identify DEGs are outlined in Table 2, categorized by the disorder and corresponding study. These methods included RNA-seq, single-nucleus RNA-seq (snRNA-seq), Bulk RNA-seq, Microarray, Robust Multi-array Average (RMA), small RNA-seq, microRNA (miRNA) expression, single-cell RNA sequencing (scRNA-seq), next-generation sequencing (NGS), mRNA sequencing, quantitative PCR (qPCR), and complementary DNA (cDNA) microarray.

### Statistical analysis

When the FDR was not calculated in the original study, we used the Benjamini-Hochberg procedure to compute the q-value to control the FDR. DEGs with FDR<0.05 were considered for the analysis. If a study included transcriptomic data from both patients and animal models, we exclusively utilized the transcriptomic data from the patients.

Custom-written MATLAB scripts were used for performing a statistical analysis of the data (see Code availability). First, the script went over all gene symbols detected as DEGs and checked with the Human Gene Nomenclature Committee (HGNC) archive whether the gene symbols are the most updated ones, to create consistency among the data. If a gene symbol was detected as previously approved by the HGNC or as an alias name (another symbol name that is used to refer to this gene), it was converted to the HGNC-approved gene symbol. Then, the script organized the identified DEGs into an Excel file named DEGs summary (Supplementary Table S5 – DEGs summary raw data). Rows in the DEGs Summary file represent a unique gene symbol, while columns represent a unique neurological disorder. The number in each position in the file represents the number of studies in which the gene was identified to be differently expressed between the neurological patients and the healthy control group. The data was normalized by the total number of studies analyzed for a specific disorder (Supplementary Table S5 – normalized DEGs summary) with the following equation. -

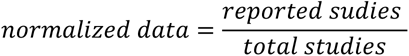

Where normalized data is the fraction of studies in which a gene was reported to be differentially expressed for a specific disorder. Reported studies is the number of studies in which a gene was reported to be differentially expressed for a specific disorder, and the total studies is the number of studies analyzed for a specific disorder. Measuring the percentage of overlapping genes between different datasets is a common method of representing the reproducibility of gene lists [70]. We used Seaborn, a Python data visualization library to create a heatmap showing the fraction of studies in which genes were reported to be differentially expressed.

### Gene Ontology (GO) analysis

To seek common dysregulated pathways, the enrichment for biological processes, cellular components, molecular functions, and KEGG was tested using ShinyGO 0.77 with default settings [1]. We used the STRING database for protein-protein interactions (PPI) and network enrichment to show interactions between dysregulated genes [2].

### Code availability

The MATLAB code for running the main analyses is available from the authors upon a reasonable request.

## Discussion

Our primary discovery involves the identification of 31 significant DEGs (FDR<0.05) shared between all seven neurological disorders compared to healthy control individuals. These DEGs included SST, BCL6, PRMT8, NFIX, COL1A2, SAT1, MT1E, SPON1, PODXL, CCN2, HLA-E, CRH, ADAMTS9, PLP1, LRRN2, TM4SF1, and more. These DEGs demonstrate significant enrichment for ECM pathways, External encapsulating structure, Response to acid chemical, Blood vessel development, Cell migration, Hormon-mediated apoptotic signalling pathway, and Receptor activity (Fig. 8A). Remarkedly, dysregulation of ECM was presented in all three categories of disorders: neurodegenerative diseases (Fig. 8B), neuropsychiatric disorders (Fig. 8C), and neurodevelopmental disorders (Fig. 8D), and also in the assessment of common pathways between ASD and Schizophrenia (Fig 9G). Furthermore, we identified the shared DEGs and associated dysregulated pathways for each category of disorders (Fig 8B, C, D), and individually for each disorder (Fig 9A-F). Finally, to increase the reproducibility, we pinpointed the most prominent DEGs (FDR<0.05) according to the fraction of reportings, that are consistently reported within and across examined disorders (not necessarily shared among all of them), these included GFAP, IFITM3, HSPA1A, MKNK2, ITGB4, SST, and more (Fig 9H).

**Figure 8.**
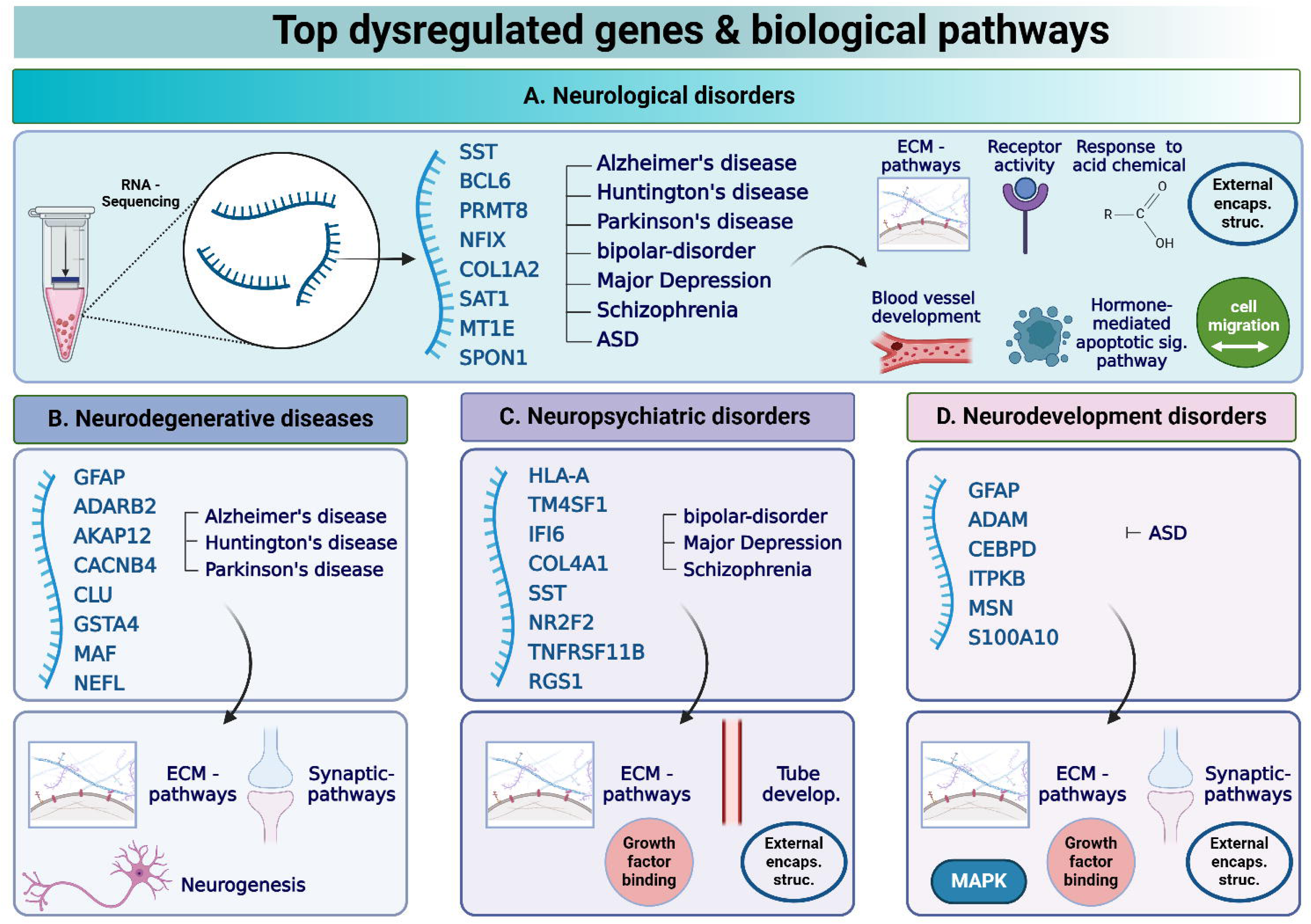
Top shared dysregulated genes and biological pathways for Neurological disorders, and per each category of disorders. **A**. A summary of the most prominent dysregulated genes and associated pathways shared among seven neurological disorders. **B**. A summary of the most prominent dysregulated genes and associated pathways shared among three neurodegenerative disease. **C**. A summary of the most prominent dysregulated genes and associated pathways shared among three Neuropsychiatric disorders. **D**. A summary of the most prominent dysregulated genes and associated pathways shared among Neurodevelopmental disorders. Abbreviations: ECM – Extracellular Matrix, External enacps. Struc – External encapsulating structure, Tube develop. – Tube development. The figure was Created with BioRender.com.

**Figure 9.**
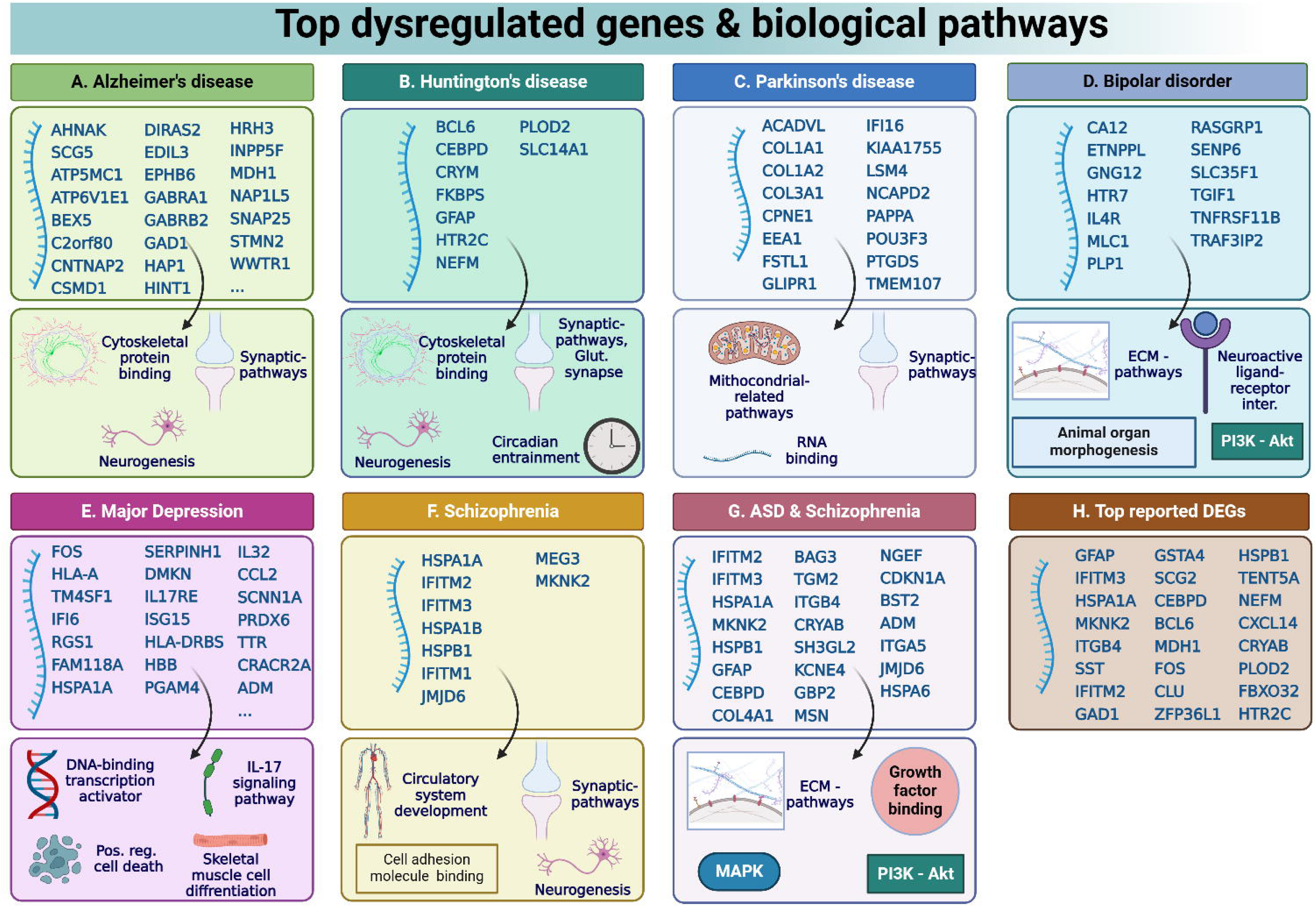
Top Dysregulated genes and biological pathways per disorder, and for transcriptional commonalities between ASD and Schizophrenia. **A**. The figure summarizes the most frequently reported dysregulated genes and associated pathways for Alzheimer’s disease. **B**. The figure summarizes the most frequently reported dysregulated genes and associated pathways for Huntington’s disease. **C**. The figure summarizes the most frequently reported dysregulated genes and associated pathways for Parkinson’s disease. **D**. The figure summarizes the most frequently reported dysregulated genes and associated pathways for bipolar disorder. **E**. The figure summarizes the most frequently reported dysregulated genes and associated pathways for Major Depression. **F**. The figure summarizes the most frequently reported dysregulated genes and associated pathways for Schizophrenia. **G**. The figure summarizes the most frequently reported dysregulated genes and associated pathways for transcriptional commonalities between ASD and Schizophrenia. **H.** The figure shows the DEGs that were reported the highest number of times among the seven disorders (not necessarily shared among all of them). GFAP, followed by IFITM3, were found to be the most reported genes. Abbreviations: ASD - Autism spectrum disorder, ECM – Extracellular Matrix, Glut synapse – Glutamatergic synapse, Neuroactive ligand-receptor Inter. – Neuroactive ligand-receptor Interaction, Pos. reg. cell death – post regulation cell death. The figure was Created with BioRender.com.

Our analysis focused on three types of models, encompassing post-mortem brain tissues, iPSC-derived neurons, and neuronal organoids obtained from neurological patients and healthy control individuals. This approach is supported by Takahashi et al. who demonstrated the similarity of human iPSCs to human embryonic stem cells in various aspects, including morphology, proliferation, surface antigens, gene expression, epigenetic status of pluripotent cell-specific genes, and telomerase activity [71]. Additional support can be found at Chiaradia et al., which noted consistent gene expression profiles and developmental pathways in brain organoids and fetal brain samples through RNA-sequencing analysis [72]. Alternative models were deemed less suitable for our objectives. Notably, peripheral blood emerges as a crucial tool for researching the diagnosis and prognosis of diseases, including neurological disorders [73, 74]. Recently, clinical trials have commenced for blood-based biomarkers of Alzheimer’s disease. [75]. However, peripheral blood does not always accurately represent the gene expression profile of a patient’s brain. Sullivan et al. discovered approximately 50% correlation between peripheral blood and multiple brain tissues [76]. While animal models play a significant role in disease research and contribute to our understanding of molecular and biological functions in health and disease, most psychiatric disorders do not have a good animal model [77] and additionally, their gene expression profile may vary depending on the specific type of animal model examined [78].

Among the 31 DEGs (FDR<0.05) shared among the seven neurological disorders, the SST gene obtained the highest ranking, followed by BCL6, based on the average fraction of reporting among all shared DEGs. The SST gene codes for somatostatin, also known as growth hormone-inhibiting hormone (GHIH). It is a neuropeptide involved in modulating cortical circuits in the brain and cognitive function and is found in various regions of the brain. Within the cortex, it can be observed in a specific group of GABAergic neurons, serving as a marker for inhibitory interneurons. Normally, somatostatin is highly expressed throughout the brain and in the Cerebrospinal fluid (CSF). A decrease in somatostatin expression was observed in various neurological disorders, including Alzheimer’s disease, individuals with Parkinson’s disease experiencing mild or severe dementia, Huntington’s disease, bipolar disorder, Major Depression, and Schizophrenia [79]. This could result in a reduction in brain size and a disruption of neural networks and functions [79]. Rubinow et al. showed that affective disorder patients showed improvement in CSF somatostatin levels when symptoms have significantly decreased or resolved [80]. In our meta-analysis, the SST gene was highly reported in Huntington’s disease (at 80% of relevant studies), in Alzheimer’s disease (at 50% of relevant studies), and in Schizophrenia (at 43% of relevant studies).

The BCL6 gene codes for B-cell Lymphoma 6 transcription repressor and plays a crucial role in regulating the development and function of B cells. These are crucial for the immune system’s ability to recognize and respond to specific pathogens. It was also identified as an oncogene in B-cell lymphomas [81]. BCL6 was additionally recognized as playing a role in neurogenesis and serves as a crucial trigger for the transformation of neural progenitor cells (NPCs) into neurons through an epigenetic process [82]. Significantly, in our study, BCL6 was found to be differentially expressed in all 5 Huntington’s disease studies analyzed, suggesting its potential as a biomarker and therapeutic target. In Parkinson’s disease and Alzheimer’s disease, BCL6 was reported in 40% and 33% of relevant studies, respectively, and in ASD in 29% of relevant studies. However, in neuropsychiatric disorders, BCL6 was reported in only 14% to 25% of the studies.

Probably the most interesting result from our meta-analysis is the profound impact of neurological disorders on the brain ECM. Many pathways related to the brain ECM were dysregulated in all the neurological disorders that were considered. The ECM has emerged as a central focus in neuroscience [83, 84], recognized as an integral part of neural functions, especially synaptic regulation [85] [86]. Dityatev et al. introduced the concept of the ’tetra-partite synapse’ that expands the traditional understanding of synaptic structure and function. In this model, the synapse is no longer considered a simple connection between two neurons but is instead viewed as a complex network involving: pre- and post-synaptic neurons, glial cells, and the ECM [87]. The ECM constitutes a dense and dynamic network [88]. It plays a crucial role in maintaining tissue architecture and homeostasis [88], providing a microenvironment for neurons and glial cells in the CNS [89]. The ECM components which are synthesized and secreted by both neurons and glial cells are divided into three major ECM structures: the perineuronal nets (PNNs) - flexible and resilient connective structures surrounding neuronal somata, axon initial segments, dendrites, and synapses. The basement membrane (BM), also known as perivascular ECM, that enables Blood-brain barrier (BBB) integrity, with its outer part embedded in astrocytic endfeet, and the Neural interstitial matrix that enables signal transduction by binding the ECM receptors’ integrins [87] [85] [90] [91]. The ECM serves various functions throughout different stages, including development, adulthood, physiology, and pathology of the CNS [90] [85] [88]. Dysregulation of CNS ECM can result in severe conditions [88, 91]. Figure 10 illustrates the state of the CNS ECM in both health and disease [85] [88] [89] [90] [91].

**Figure 10.**
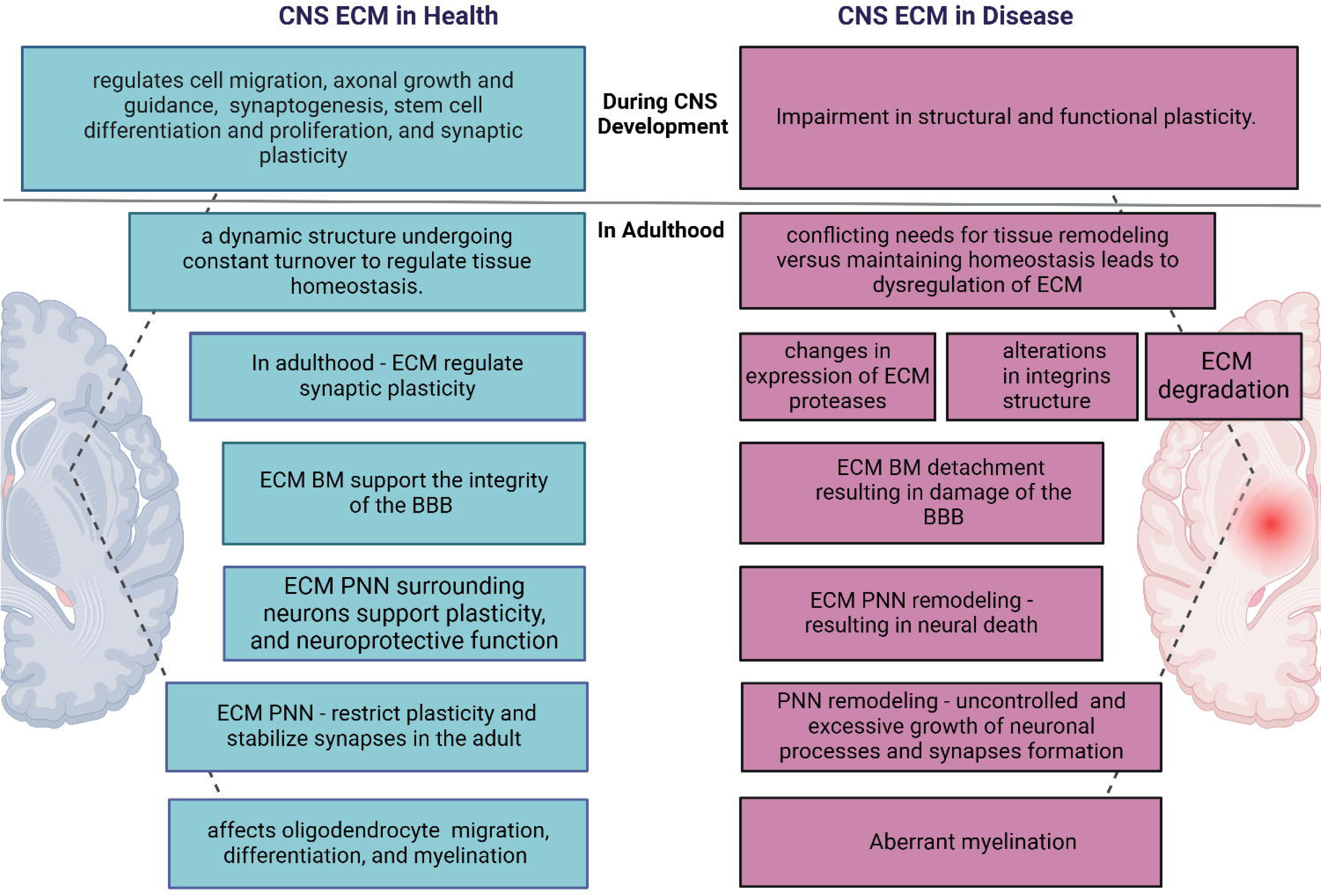
CNS ECM in health and disease. **A**. The figure illustrates the state of the CNS ECM in both health and disease during development and adulthood. Abbreviations: ECM – Extracellular Matrix, PNN - perineuronal net, BM - basement membrane, BBB - Blood-brain barrier. The figure was Created with BioRender.com.

We identified GFAP followed by IFITM3 as having the highest ranking of overlapping DEGs (FDR < 0.05) between different datasets among all disoders (Fig. 5A, Supplementary Table S1 – a list of the DEGs). Other highly reported dysregulated genes included HSPA1A, MKNK2, ITGB4, SST, IFITM2, GAD1, GSTA4, CG2, CEBPD, BCL6, MDH1, FOS, and CLU. GFAP codes for glial fibrillary acidic protein and serves as the predominant intermediate filament expressed in mature astrocytes [92]. GFAP helps to maintain the mechanical strength and shape of astrocytes that support neurons in the brain [93]. Astrocytic end-feet containing GFAP are involved in multiple functions including astrocyte motility and migration, astrocytes proliferation, and growth. They are an essential component of the blood–brain barrier (BBB), the glymphatic system [94], and have a critical role in neuron metabolic functions [95], in plasticity, and contribute to tetra-partite synapses [96]. GFAP has regulatory functions in autophagy processes, specifically, in chaperone-mediated autophagy (CMA) process [97]. Decreases in CMA function with increasing age may contribute to pathogenesis such as tau and Aβ accumulation [93]. Increased GFAP levels were presumed to result from a shift in astrocyte’s phenotype, transitioning from normal physiological astrocyte functions to reactive astrocyte functions, thereby modifying their function and disrupting the balance between neuro-supportive and neurotoxic properties [98]. In healthy individuals, CSF GFAP was demonstrated to increase with age, and is highly expressed in the aged brain [99]. Decreased levels of the GFAP gene were observed in a group of young patients with Major Depression (between 30-45 years of age) compared to healthy matched control and compared to elderly patients with Major Depression [100]. Increased levels of GFAP were observed in the superior frontal, parietal, and cerebellar cortices of ASD subjects [101], while decreased levels of GFAP were observed in frontal brain regions of Schizophrenia subjects [102]. Interestingly, GFAP was reported to be differently expressed in our meta-analysis in all examined disorders except bipolar disorder and was highly reported in neurodegenerative diseases and neurodevelopmental disorders studies.

The IFITM3 gene codes for the interferon-induced transmembrane protein 3. The IFITM3 protein functions as the first line of antiviral defense against infections [103]. Hur et al. revealed that pathogenic challenges such as aging, stroke, traumatic brain injury, or inflammatory conditions, induce the release of proinflammatory cytokines from astrocytes and microglia that could lead to an increase expression of IFITM3 in neurons and astrocytes. The IFITM3 protein binds to γ-secretase and upregulates its activity and results in increased Aβ [104]. Conversely, the accumulation of Aβ also initiates the pathology associated with Alzheimer’s disease; In their research, they observed significantly high mRNA levels of IFITM3 in post-mortem tissues in late onset Alzheimer’s disease patients (n=18) compared to non-demented control participants (n=10) [104]. We identified dysregulation of IFITM3 in 33% of Alzheimer’s disease studies. Significantly, IFITM3 was dysregulated in all neurological disorders examined in the current study except in Major Depression. IFITM3 was highly reported as differentially expressed in Huntington’s disease (80% of relevant studies), in Schizophrenia (71% of relevant studies), and in Parkinson’s disease (60% of relevant studies).

Among the shared DEGs between both ASD and Schizophrenia, IFITM2 emerged as the most frequently reported gene, followed by HSPB1, IFITM3, HSPA1A, MKNK2, GFAP, COL4A1, TGM2, BAG3, and CEBPD. Enrichment analysis revealed significant associations with ECM pathways in terms of cellular components, biological processes, and molecular functions. Notably, the HSPB1 gene, encoding for heat shock protein family B (small) member 1, also known as HSP27, contributes significantly to cell survival in stressful conditions. In addition, HSPB1 has an important role during CNS development. This aligns with the neurodevelopmental hypothesis of Schizophrenia, suggesting abnormal or disrupted neural growth during embryonic development [27]. In our meta analysis, HSPB1 was identified in 57.1% of both, ASD and Schizophrenia studies.

The central aspects of our findings highlight shared features within and across neurological disorders, underscoring a notable distinction in the ECM of the CNS between health and disease.Understanding these differences is crucial for pinpointing potential biomarkers and developing therapeutic strategies that target the CNS ECM to promote tissue repair and recovery in neurological disorders. The shared features among disorders may also indicate overlap in symptoms and comorbidity between brain disorders. Further investigations are essential, providing detailed insights into gene expression levels, whether they undergo upregulation or downregulation, as well as DNA and protein level verification. These examinations are imperative to elucidate the precise roles of the identified lead genes. Additionally, exploring the relationships among these leading genes is warranted.

---

| <u>Disease</u> | <u>Article's Title</u> | <u>model</u> | <u>No.of DEG</u><br><u>FDR<0.05</u><br><u>Included</u> | <u>area in the brain</u> | <u>transcriptomic analysis</u> | <u>patient</u> | <u>control</u> | <u>Authors</u> | <u>Year Published</u> |
| --- | --- | --- | --- | --- | --- | --- | --- | --- | --- |
| Alzheimer | Single-cell transcriptomic analysis of Alzheimer's disease | post-mortem tissue | 2238 | prefrontal cortex (BA 10) | single-nucleus RNA-seq (snRNA-seq) , Bulk RNA-seq | 24 | 24 | Mathys et al. | 2019 |
| Alzheimer | A single-cell atlas of entorhinal cortex from individuals with Alzheimer's disease reveals cell-type-specific gene expression regulation | post-mortem tissue | 459 | entorhinal cortex | single-nucleus RNA sequencing (DroNc-Seq) | 6 | 6 | Grubman et al. | 2019 |
| Alzheimer | APOE4 exacerbates synapse loss and neurodegeneration in Alzheimer's disease patient iPSC-derived cerebral organoids | iPSC organoids | 1265 | 3-D cerebral organoid models using human iPSC | RNA-seq | 10 | 10 | Zhao et al. | 2020 |
| Alzheimer | Defining early changes in Alzheimer's disease from RNA sequencing of brain regions differentially affected by pathology | post-mortem tissue | 589 | precuneus (PREC), primary visual cortex (VIC) | RNA-Seq | 5 | 5 | Guennewig et al. | 2021 |
| Alzheimer | Neuronal ApoE Upregulates MHC-I Expression to Drive Selective Neurodegeneration in Alzheimer's Disease | post-mortem tissue | 4063 | prefrontal cortex | snRNA-seq | 24 | 24 | A Zalocusky et al. | 2021 |
| Alzheimer | Molecular subtyping of Alzheimer's disease using RNA sequencing data reveals novel mechanisms and targets | post-mortem tissue | 1864 | frontal pole (FP), superior temporal gyrus (STG), para-hippocampal gyrus (PHG), inferior frontal gyrus (IFG) | RNA-seq | 151 | 32 | A Neff et al. | 2021 |
| ASD | Transcriptomic analysis of autistic brain reveals convergent molecular pathology | post-mortem tissue | 253 | superior temporal gyrus(BA 41/42), prefrontal cortex (BA 9) , cerebellar vermis | Microarrays and RNA-seq | 19 | 17 | Voineagu et al. | 2011 |
| ASD | NRXN1a+/- is associated with increased excitability in ASD iPSC-derived neurons | iPSC neurons | 425 | cortical neurons | RNA-seq | 3 | 5 | Avazzadeh et al. | 2021 |
| ASD | Cortical overgrowth in a preclinical forebrain organoid model of CNTNAP2-associated autism spectrum disorder | IPSC organoids | 339 | forbrain organoids, embryonic prefrontal cortical cell types | single cell RNA-Sequencing | 3 | 3 | Jong et al. | 2021 |
| ASD | Genome-wide expression assay comparison across frozen and fixed postmortem brain tissue samples | post-mortem tissue | 1567 | prefrontal cortex | Microarray | 61 | 61 | Chow et al. | 2011 |
| ASD | Brain transcriptional and epigenetic associations with autism | post-mortem tissue | 734 | cerebellar, Occipital lobe (BA 19) | Microarray | 9 | 9 | Ginsberg et al. | 2012 |
| ASD | Immune transcriptome alterations in the temporal cortex of subjects with autism | post-mortem tissue | 138 | superior temporal gyrus | Microarray | 6 | 6 | Garbett et al. | 2008 |
| ASD | IQSEC2 mutation associated with epilepsy, intellectual disability, and autism results in hyperexcitability of patient-derived neurons and deficient synaptic transmission | IPSC neurons | 4001 | DG granule neurons | RNA sequencing | 1 | 1 | Brant et al. | 2021 |
| bipolar | Transcriptome Sequencing of the Anterior Cingulate in Bipolar Disorder: Dysregulation of G Protein-Coupled Receptors | post-mortem tissue | 10 | anterior cingulate cortex (ACC) (BA 24) | RNA sequencing | 13 | 13 | Cruceanu et al. | 2015 |
| bipolar | Altered expression of mitochondria-related genes in postmortem brains of patients with bipolar disorder or schizophrenia, as revealed by large-scale DNA microarray analysis | post-mortem tissue | 5 | prefrontal cortex (BA 46) | qRT-PCR and microarray | 33 | 34 | Iwamoto et al. | 2004 |
| bipolar | Oligodendrocyte dysfunction in schizophrenia and bipolar disorder | post-mortem tissue | 9 | prefrontal cortex (BA 9) | qPCR and Microarray | 15 | 15 | Tkachev et al. | 2003 |
| bipolar | Coding and Non-Coding RNA Abnormalities in | post-mortem tissue | 31 | medial frontal gyrus | RNA sequencing | 4 | 4 | Luykx et al. | 2019 |
| bipolar | Deficient *LEF1* expression is associated with lithium resistance and hyperexcitability in neurons derived from bipolar disorder patients | IPSC neurons | 817 | hippocampal dentate gyrus-like neurons | RNA-seq analysis | 6 | 4 | Santos et al. | 2021 |
| bipolar | Gene expression analysis of bipolar disorder reveals downregulation of the ubiquitin cycle and alterations in synaptic genes | post-mortem tissue | 78 | prefrontal cortex (BA 9), orbitofrontal cortex (BA 11) | microarray technology | 10 | 11 | Ryan et al. | 2006 |
| bipolar | A combined analysis of genome-wide expression profiling of bipolar disorder in human prefrontal cortex | post-mortem tissue | 62 | prefrontal cortex (BA 9, 10, 11 or 46) | Robust Multi-array Average (RMA) | 101 | 106 | Wang et al. | 2016 |
| Huntington | Single-nucleus RNA-seq identifies Huntington disease astrocyte states | post-mortem tissue | 4494 | anterior cingulate cortex (BA 24) | 4 snRNAseq & 12 bulk RNAseq | 8 | 8 | Dalahmah et al. | 2020 |
| Huntington | Transcriptome sequencing reveals aberrant alternative splicing in Huntington's disease | post-mortem tissue | 92 | primary motor cortex (BA 4) | RNA-seq analysis | 7 | 7 | Lin et al. | 2016 |
| Huntington | Huntington's disease-specific mis-splicing unveils key effector genes and altered splicing factors | post-mortem tissue | 7412 | striatum | RNA-seq analysis | 3 | 3 | Elorza et al. | 2021 |
| Huntington | Regional and cellular gene expression changes in human Huntington's disease brain | post-mortem tissue | 8737 | Cerebellar, Caudate, Frontal cortex | microarrays | 44 | 36 | Hodges et al. | 2006 |
| Huntington | Cell Type-Specific Transcriptomics Reveals that Mutant Huntingtin Leads to Mitochondrial RNA Release and Neuronal Innate Immune Activation | post-mortem tissue | 4161 | caudate, putamen | snRNA-Seq | 2-4 | 2-4 | Lee et al. | 2020 |
| Major Depression | Altered neuro-inflammatory gene expression in hippocampus in major depressive disorder | post-mortem tissue | 162 | hippocampus (right temporal lobe) | RNA-seq analysis | 23 | 23 | Mahajan et al. | 2018 |
| Major Depression | Sex-specific transcriptional signatures in human depression | post-mortem tissue | 12099 | ventromedial PFC, OFC, DLPFC, anterior insula, nucleus accumbens, ventral subiculum | RNA-seq analysis | 26 | 22 | Labonté et al. | 2017 |
| Major Depression | Whole-transcriptome brain expression and exon-usage profiling in major depression and suicide: evidence for altered glial, endothelial and ATPase activity | post-mortem tissue | 9 | dorsal lateral prefrontal cortex (DLPFC) (BA 9) | small RNA-seq, miRNA expression | 30 | 29 | Pantazatos et al. | 2017 |
| Major Depression | Altered expression of genes involved in inflammation and apoptosis in frontal cortex in major depression | post-mortem tissue | 9 | prefrontal cortex (BA10) | microarray mRNA expression analysis | 14 | 14 | Shelton et al. | 2011 |
| Parkinson | Differential transcript usage in the Parkinson's disease brain | post-mortem tissue | 599 | prefrontal cortex (BA 9) | RNA-Seq (DEXSeq and DRIMSeq) | 27 | 22 | Dick et al. | 2020 |
| Parkinson | The Parkinson's-disease-associated mutation LRRK2-G2019S alters dopaminergic differentiation dynamics via NR2F1 | IPSC neurons | 2389 | dopaminergic (DA) neurons | scRNA-seq | 3 | 4 | Walter et al. | 2021 |
| Parkinson | Reduced synaptic activity and dysregulated extracellular matrix pathways in midbrain neurons from Parkinson's disease patients | IPSC neurons | 6414 | dopaminergic (DA) neurons | RNA-seq analysis | 27 | 20 | stern et al. | 2022 |
| Parkinson | Upregulated extracellular matrix-related genes and impaired synaptic activity in dopaminergic and hippocampal neurons derived from Parkinson's disease patients with *PINK1* and *PARK2* mutations | IPSC neurons | 2601 | Midbrain - Dopaminergic (DA) neurons, Hippocampal - DG (Dentate gyrus) granule neurons | RNA-seq analysis | 2 | 2 | Tripathi et al. | 2023 |
| Parkinson | Synaptic dysfunction and extracellular matrix dysregulation in dopaminergic neurons from sporadic and E326K-GBA1 Parkinson's disease patients | IPSC neurons | 1435 | midbrain DA neurons | next-generation sequencing (NGS) | 6 | 3 | Rosh et al. | 2024 |
| Schizophrenia | Non-coding RNA dysregulation in the amygdala region of schizophrenia patients contributes to the pathogenesis of the disease | post-mortem tissue | 1182 | amygdala, DLPFC | RNA-seq | 22 | 24 | Liu et al. | 2018 |
| Schizophrenia | RNA-seq analysis of amygdala tissue reveals characteristic expression profiles in schizophrenia | post-mortem tissue | 303 | amygdala, hippocampus | RNA-seq | 22 | 24 | Chang et al. |  |
| Schizophrenia | Gene expression profiling by mRNA sequencing reveals increased expression of immune/inflammation-related genes in the hippocampus of individuals with schizophrenia | post-mortem tissue | 144 | hippocampus | RNA-Seq | 14 | 15 | Hwang et al. | 2013 |
| Schizophrenia | Transcriptome sequencing revealed significant alteration of cortical promoter usage and splicing in schizophrenia | post-mortem tissue | 772 | ortical grey matter of STG from the left hemisphere | mRNA sequencing | 9 | 9 | Wu et al. | 2012 |
| Schizophrenia | Migratory cortical interneuron-specific transcriptome abnormalities in schizophrenia | IPSC neurons | 96 | Cortical interneurons (cINs) | RNA-seq analysis | 5 | 5 | Park et al. | 2021 |
| Schizophrenia | Modelling schizophrenia using human induced pluripotent stem cells | IPSC neurons | 589 | glutamatergic neurons | Microarray & qPCR | 4 | 6 | Brennand et al. | 2011 |
| Schizophrenia | Analysis of gene expression in two large schizophrenia cohorts identifies multiple changes associated with nerve terminal function | post-mortem tissue | 1521 | anterior prefrontal cortex (BA10) | Microarray | 28 | 23 | Maycox et al. | 2009 |

