## Extended figure 1 for "Brain Extracellular Matrix implications in multiple neurological disorders are revealed through a meta-analysis of transcriptional changes"

### A Alzheimer's disease Top DEGs

| Total No. of Alzheimer's disease studies analyzed: 6 | | |
|---|---|---|
| **Gene Symbol** | **Name** | **Percentage of Alzheimer's diseases studies reported these DEGs** |
| AHNAK | AHNAK nucleoprotein | 83.3% |
| SCG5 | secretogranin V | 83.3% |
| ATP5MC1 | ATP synthase membrane subunit c locus 1 | 66.7% |
| ATP6V1E1 | ATPase H+ transporting V1 subunit E1 | 66.7% |
| BEX5 | brain expressed X-linked 5 | 66.7% |
| C2orf80 | chromosome 2 open reading frame 80 | 66.7% |
| CNTNAP2 | contactin associated protein 2 | 66.7% |
| CSMD1 | CUB and Sushi multiple domains 1 | 66.7% |
| DIRAS2 | DIRAS family GTPase 2 | 66.7% |
| EDIL3 | EGF like repeats and discoidin domains 3 | 66.7% |
| EPHB6 | EPH receptor B6 | 66.7% |
| GABRA1 | gamma-aminobutyric acid type A receptor subunit alpha1 | 66.7% |
| GABRB2 | gamma-aminobutyric acid type A receptor subunit beta2 | 66.7% |
| GAD1 | glutamate decarboxylase 1 | 66.7% |
| HAP1 | huntingtin associated protein 1 | 66.7% |
| HINT1 | histidine triad nucleotide binding protein 1 | 66.7% |
| HRH3 | histamine receptor H3 | 66.7% |
| INPP5F | inositol polyphosphate-5-phosphatase F | 66.7% |
| ITPKB | inositol-trisphosphate 3-kinase B | 66.7% |
| JPH3 | junctophilin 3 | 66.7% |
| KCNIP4 | potassium voltage-gated channel interacting protein 4 | 66.7% |
| KCNN3 | potassium calcium-activated channel subfamily N membe | 66.7% |
| LRRTM4 | leucine rich repeat transmembrane neuronal 4 | 66.7% |
| MDH1 | malate dehydrogenase 1 | 66.7% |
| MEF2C | myocyte enhancer factor 2C | 66.7% |
| MTSS2 | MTSS I-BAR domain containing 2 | 66.7% |
| NAP1L5 | nucleosome assembly protein 1 like 5 | 66.7% |
| NAPB | NSF attachment protein beta | 66.7% |
| NCS1 | neuronal calcium sensor 1 | 66.7% |
| NRN1 | neuritin 1 | 66.7% |
| OPCML | opioid binding protein/cell adhesion molecule like | 66.7% |
| PLD1 | phospholipase D1 | 66.7% |
| PLEC | plectin | 66.7% |
| PLXNB1 | plexin B1 | 66.7% |
| POLR2K | RNA polymerase II, I and III subunit K | 66.7% |
| RAB3A | RAB3A, member RAS oncogene family | 66.7% |
| RGS4 | regulator of G protein signaling 4 | 66.7% |
| RIN3 | Ras and Rab interactor 3 | 66.7% |
| RPH3A | rabphilin 3A | 66.7% |
| SNAP25 | synaptosome associated protein 25 | 66.7% |
| SNX10 | sorting nexin 10 | 66.7% |
| STMN2 | stathmin 2 | 66.7% |
| TGFB1 | transforming growth factor beta 1 | 66.7% |
| VDAC1 | voltage dependent anion channel 1 | 66.7% |
| VSNL1 | visinin like 1 | 66.7% |
| WWTR1 | WW domain containing transcription regulator 1 | 66.7% |

### B Cellular Components

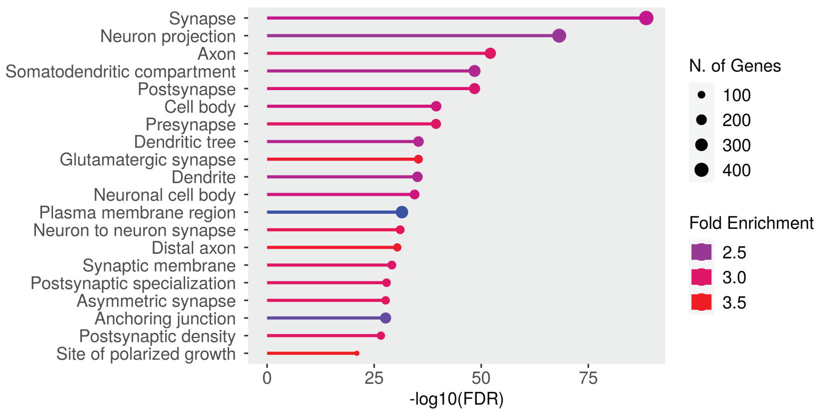


### C Biological Processes

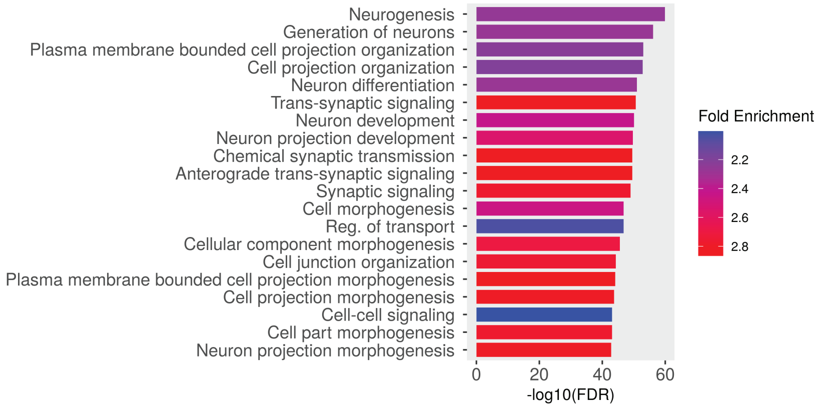


### D Molecular Function

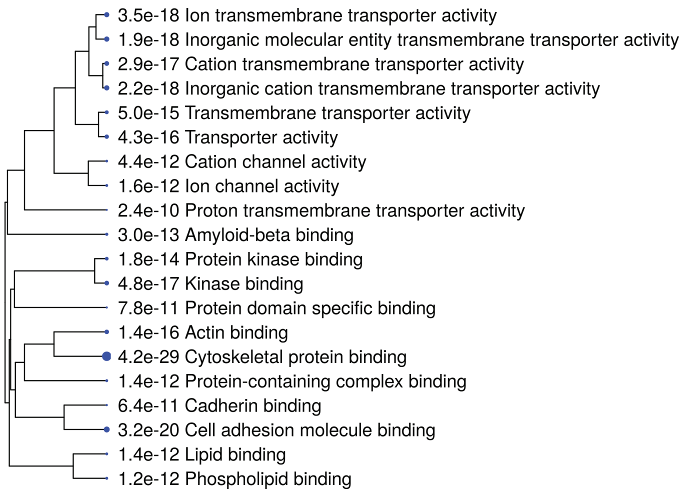


### E KEGG

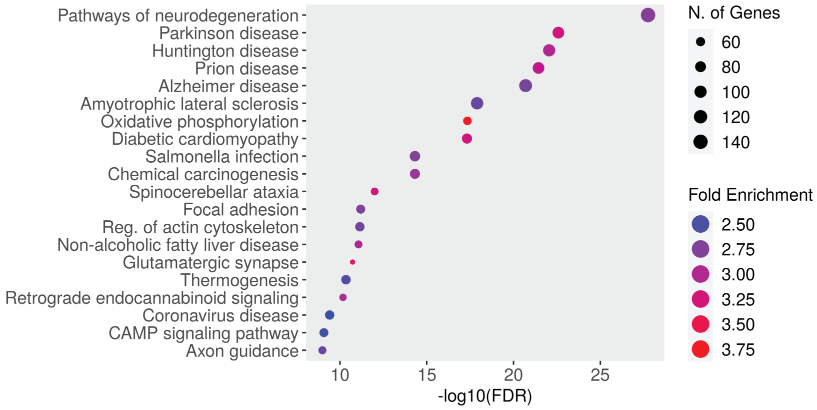


### F Gene Type

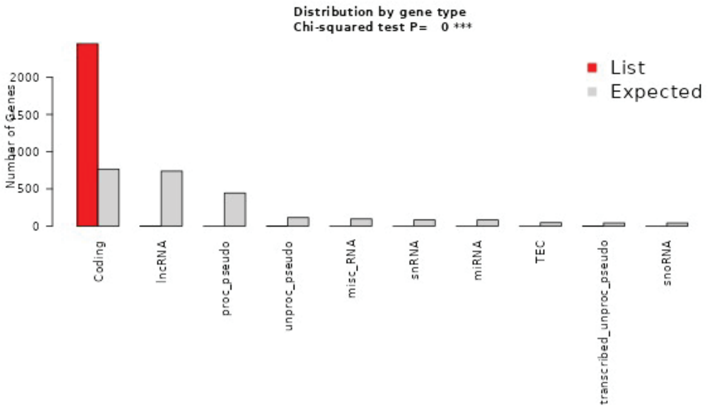
