## Extended figure 2 for "Brain Extracellular Matrix implications in multiple neurological disorders are revealed through a meta-analysis of transcriptional changes"

### A Parkinson's disease Top DEGs

| Total No. of Parkinson's disease studies analyzed: 5 | | |
|---|---|---|
| **Gene Symbol** | **Name** | **Percentage of Parkinson's diseases studies reported these DEGs** |
| ACADVL | acyl-CoA dehydrogenase very long chain | 80.0% |
| COL1A1 | collagen type I alpha 1 chain | 80.0% |
| COL1A2 | collagen type I alpha 2 chain | 80.0% |
| COL3A1 | collagen type III alpha 1 chain | 80.0% |
| CPNE1 | copine 1 | 80.0% |
| EEA1 | early endosome antigen 1 | 80.0% |
| FSTL1 | follistatin like 1 | 80.0% |
| GLIPR1 | GLI pathogenesis related 1 | 80.0% |
| IFI16 | interferon gamma inducible protein 16 | 80.0% |
| KIAA1755 | KIAA1755 | 80.0% |
| LSM4 | LSM4 homolog, U6 small nuclear RNA and mRNA degradation associate | 80.0% |
| NCAPD2 | non-SMC condensin I complex subunit D2 | 80.0% |
| PAPPA | pappalysin 1 | 80.0% |
| POU3F3 | POU class 3 homeobox 3 | 80.0% |
| PTGDS | prostaglandin D2 synthase | 80.0% |
| TMEM107 | transmembrane protein 107 | 80.0% |

### B Cellular Components

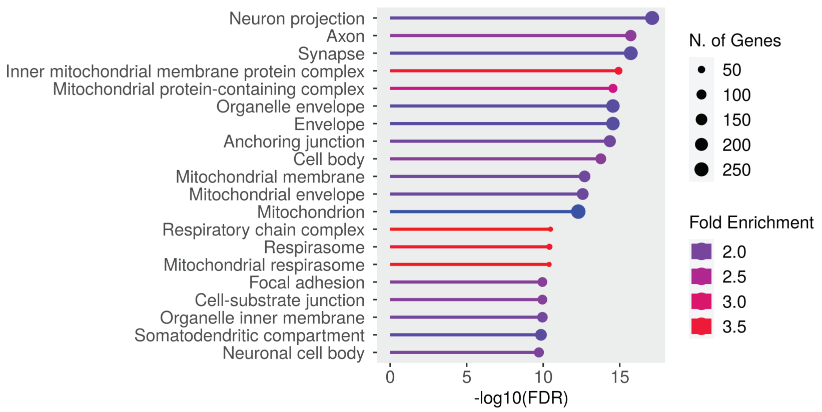


### C Biological Processes

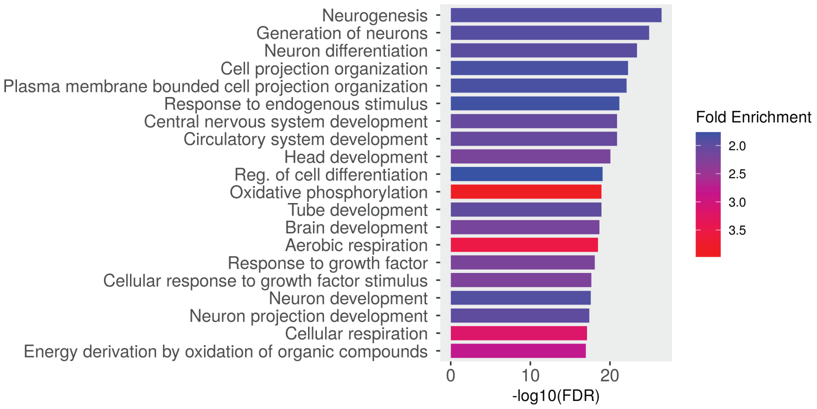


### D Molecular Function

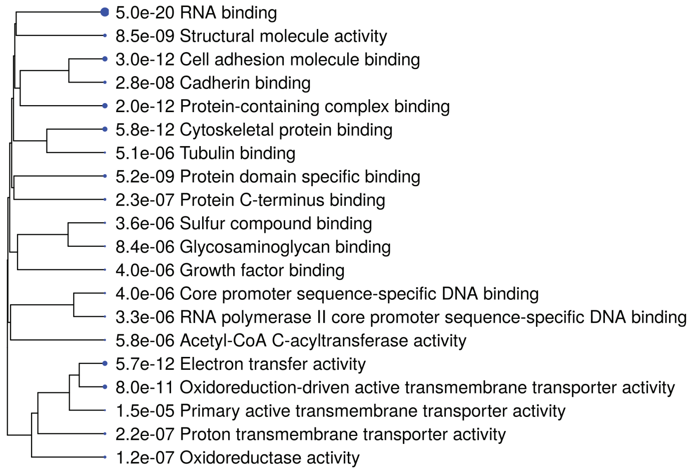


### E KEGG

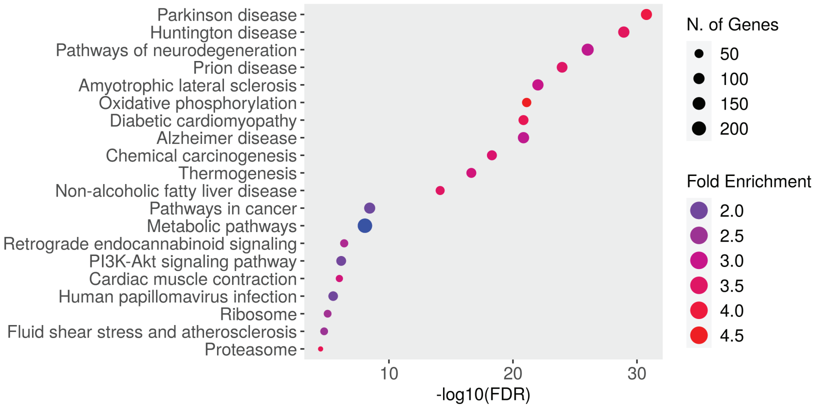


### F Gene Type

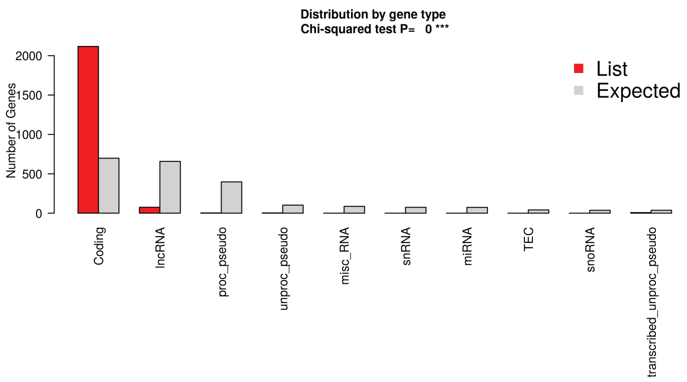
