## Extended figure 3 for "Brain Extracellular Matrix implications in multiple neurological disorders are revealed through a meta-analysis of transcriptional changes"

### A Huntington's disease Top DEGs

| Total No. of Huntington's disease studies analyzed:5 | | |
|---|---|---|
| **Gene Symbol** | **Name** | **Percentage of Huntington's diseases studies reported these DEGs** |
| BCL6 | BCL6 transcription repressor | 100.0% |
| CEBPD | CCAAT enhancer binding protein delta | 100.0% |
| CRYM | crystallin mu | 100.0% |
| FKBP5 | FKBP prolyl isomerase 5 | 100.0% |
| GFAP | glial fibrillary acidic protein | 100.0% |
| HTR2C | 5-hydroxytryptamine receptor 2C | 100.0% |
| NEFM | neurofilament medium chain | 100.0% |
| PLOD2 | procollagen-lysine,2-oxoglutarate 5-dioxygenase | 100.0% |
| SLC14A1 | solute carrier family 14 member 1 (Kidd blood grou | 100.0% |

### B top reported DEGs - PPIs networks

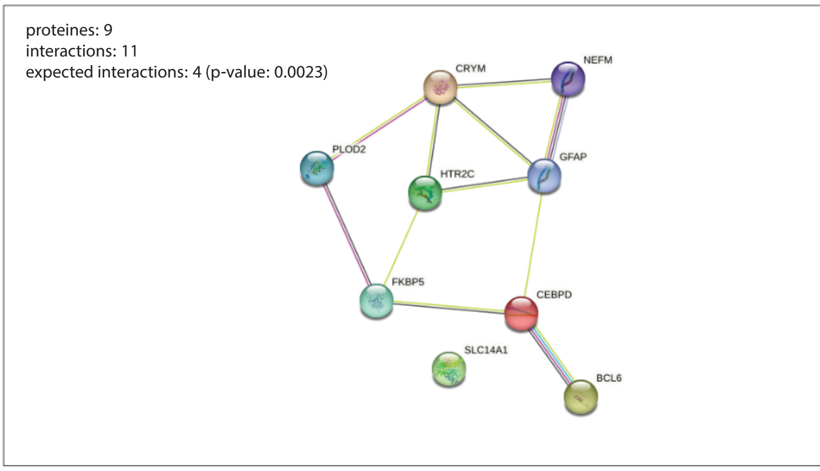


### C Gene Type

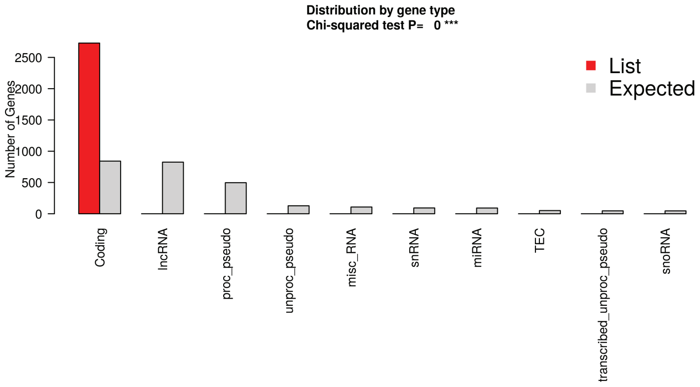


### D Cellular Components

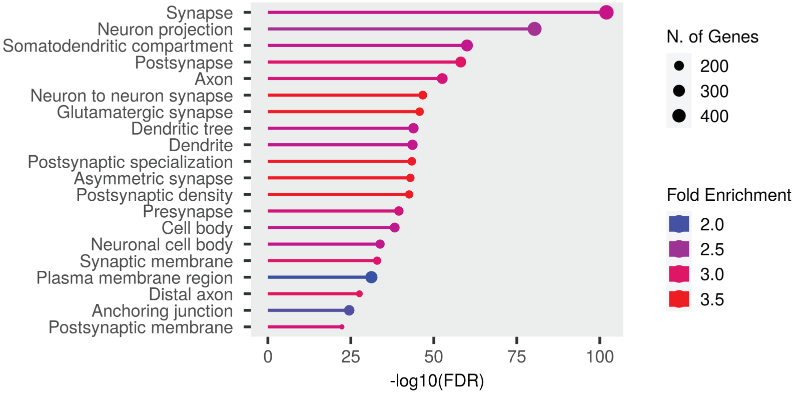


### E Biological Processes

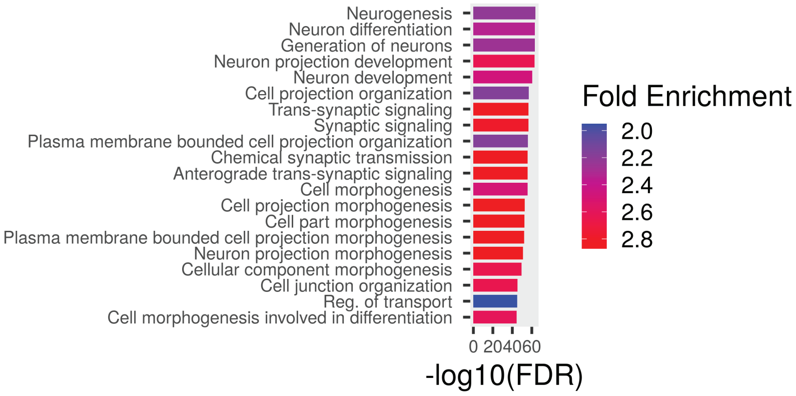


### F Molecular Function

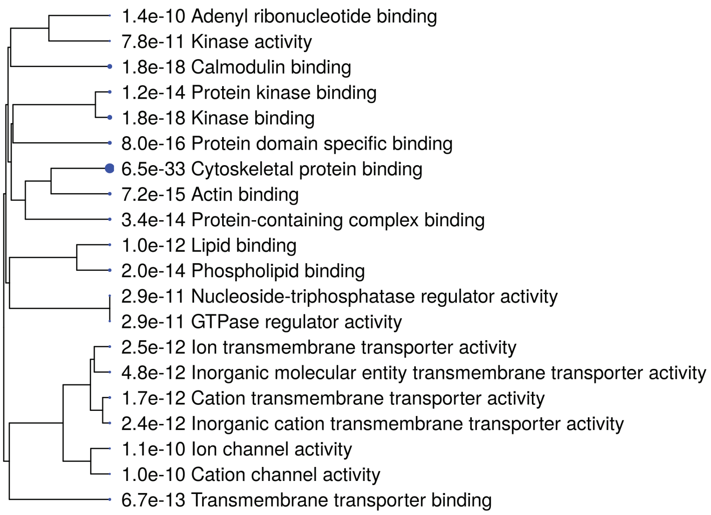


### G KEGG

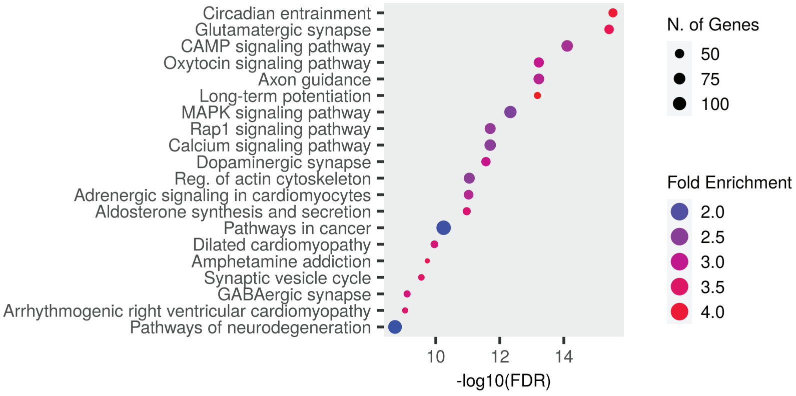
