## Extended figure 4 for "Brain Extracellular Matrix implications in multiple neurological disorders are revealed through a meta-analysis of transcriptional changes"

A Schizophrenia Top reported DEGs

| Total No. of Schizophrenia's studies analyzed: 7 | | |
|---|---|---|
| **gene_symbol** | **name** | **Percentage of Schizophrenia's studies reported these DEGs** |
| HSPA1A | heat shock protein family A (Hsp70) member 1A | 71.4% |
| IFITM3 | interferon induced transmembrane protein 3 | 71.4% |
| IFITM2 | interferon induced transmembrane protein 2 | 71.4% |
| MKNK2 | MAPK interacting serine/threonine kinase 2 | 57.1% |
| HSPA1B | heat shock protein family A (Hsp70) member 1B | 57.1% |
| MEG3 | maternally expressed 3 | 57.1% |
| HSPB1 | heat shock protein family B (small) member 1 | 57.1% |
| JMJD6 | jumonji domain containing 6, arginine demethylase | 57.1% |
| IFITM1 | interferon induced transmembrane protein 1 | 57.1% |

B Cellular Components

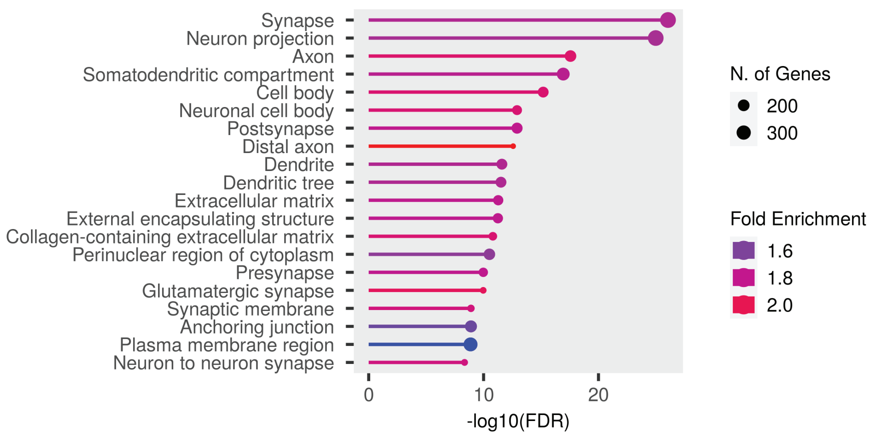


C Biological Processes

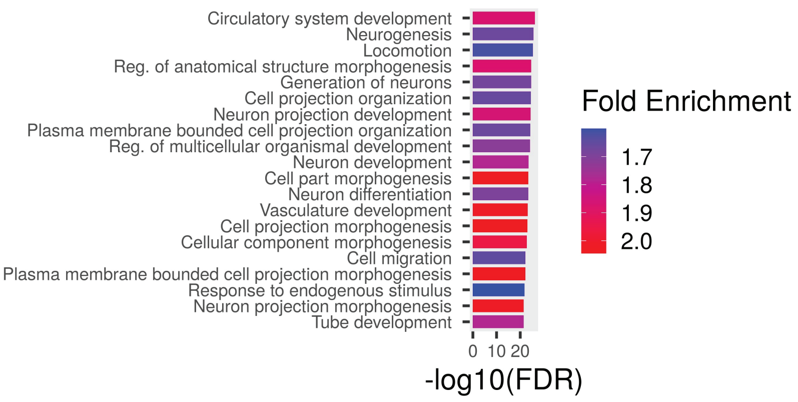


D Molecular Function

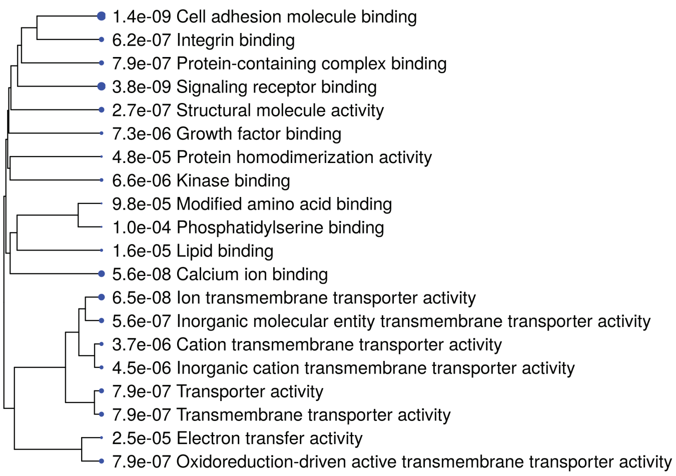


E KEGG

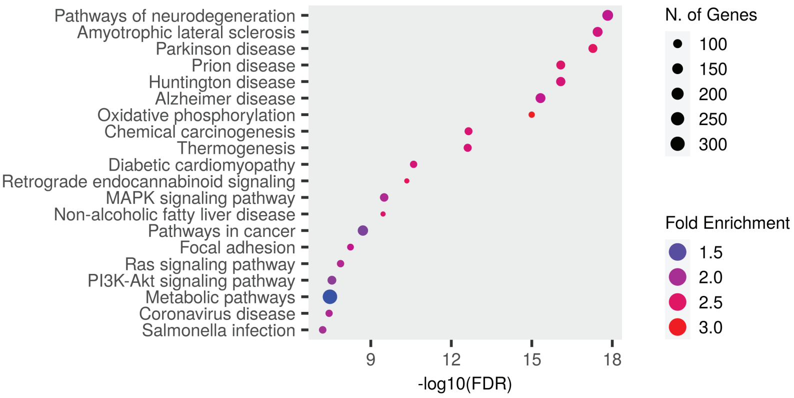


F Gene Type

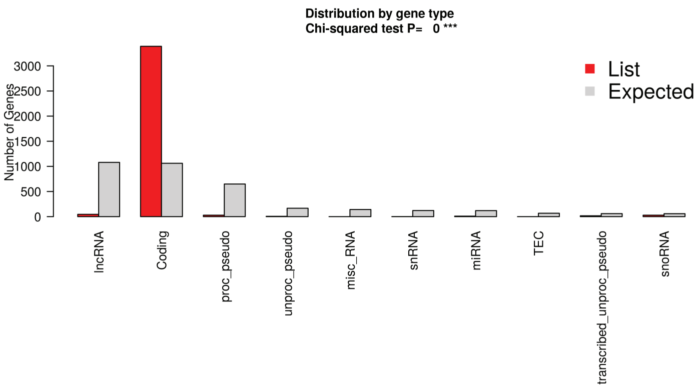
