## Extended figure 5 for "Brain Extracellular Matrix implications in multiple neurological disorders are revealed through a meta-analysis of transcriptional changes"

A Major Depression - Top reported DEGs

| Total No. of Major Depression's studies analyzed: 4 | | |
|---|---|---|
| gene_symbol | name | Percentage of Major Depression's studies reported these DEGs |
| FOS | Fos proto-oncogene, AP-1 transcription factor subu | 75.0% |
| ADM | adrenomedullin | 50.0% |
| ATF3 | activating transcription factor 3 | 50.0% |
| CCL2 | C-C motif chemokine ligand 2 | 50.0% |
| CD84 | CD84 molecule | 50.0% |
| CRACR2A | calcium release activated channel regulator 2A | 50.0% |
| DDHD1 | DDHD domain containing 1 | 50.0% |
| DMKN | dermokine | 50.0% |
| DUSP1 | dual specificity phosphatase 1 | 50.0% |
| EGR1 | early growth response 1 | 50.0% |
| FAM118A | family with sequence similarity 118 member A | 50.0% |
| HBB | hemoglobin subunit beta | 50.0% |
| HLA-A | major histocompatibility complex, class I, A | 50.0% |
| HLA-DRB1 | major histocompatibility complex, class II, DR beta | 50.0% |
| HLA-DRB5 | major histocompatibility complex, class II, DR beta | 50.0% |
| HSPA1A | heat shock protein family A (Hsp70) member 1A | 50.0% |
| IFI6 | interferon alpha inducible protein 6 | 50.0% |
| IL17RE | interleukin 17 receptor E | 50.0% |
| IL32 | interleukin 32 | 50.0% |
| ISG15 | ISG15 ubiquitin like modifier | 50.0% |
| KCNQ1OT1 | KCNQ1 opposite strand/antisense transcript 1 | 50.0% |
| MBTPS2 | membrane bound transcription factor peptidase, si | 50.0% |
| MIR663A | microRNA 663a | 50.0% |
| MTPAP | mitochondrial poly(A) polymerase | 50.0% |
| NPAS4 | neuronal PAS domain protein 4 | 50.0% |
| NR4A1 | nuclear receptor subfamily 4 group A member 1 | 50.0% |
| PGAM4 | phosphoglycerate mutase family member 4 | 50.0% |
| POLR1H | RNA polymerase I subunit H | 50.0% |
| PRDX6 | peroxiredoxin 6 | 50.0% |
| PRKY | protein kinase Y-linked (pseudogene) | 50.0% |
| PVT1 | Pvt1 oncogene | 50.0% |
| RGS1 | regulator of G protein signaling 1 | 50.0% |
| RGS17 | regulator of G protein signaling 17 | 50.0% |
| RNY1 | RNA, Ro60-associated Y1 | 50.0% |
| RPL7A | ribosomal protein L7a | 50.0% |
| SCNN1A | sodium channel epithelial 1 subunit alpha | 50.0% |
| SERPINH1 | serpin family H member 1 | 50.0% |
| SLC12A8 | solute carrier family 12 member 8 | 50.0% |
| SNHG29 | small nucleolar RNA host gene 29 | 50.0% |
| TM4SF1 | transmembrane 4 L six family member 1 | 50.0% |
| TTR | transthyretin | 50.0% |
| XIST | X inactive specific transcript | 50.0% |
| y_rna | small non-coding RNAs | 50.0% |

B Cellular Components - High level Category

| No. of genes | High level GO category | Genes Symbol |
|---|---|---|
| 14 | Extracellular region | IL32 CCL2 SCNN1A PRDX6 TTR CRACR2A ADM SERPINH1 DMKN IL17RE ISG15 HLA-DRB5 PGAM4 HBB |
| 11 | Extracellular space | IL32 CCL2 SCNN1A PRDX6 TTR ADM SERPINH1 DMKN HLA-DRB5 PGAM4 HBB |
| 6 | Organelle membrane | MBTPS2 SCNN1A NR4A1 IFI6 CRACR2A HLA-DRB5 |
| 6 | Extracellular organelle | SCNN1A PRDX6 TTR HLA-DRB5 PGAM4 HBB |
| 5 | Chromatin | EGR1 NR4A1 ATF3 FOS NPAS4 |
| 4 | Cell projection | RGS17 SCNN1A FOS PGAM4 |

C Biological Processes

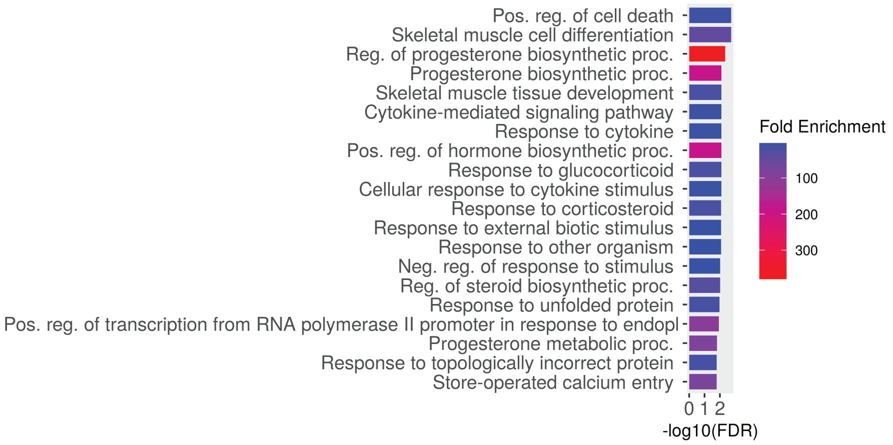


D Molecular Function

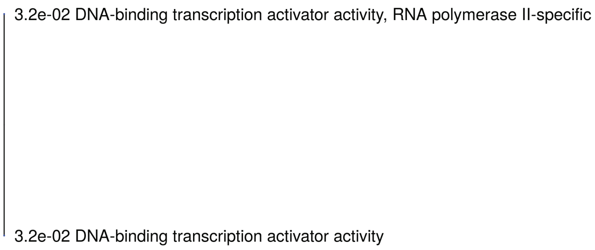


E KEGG

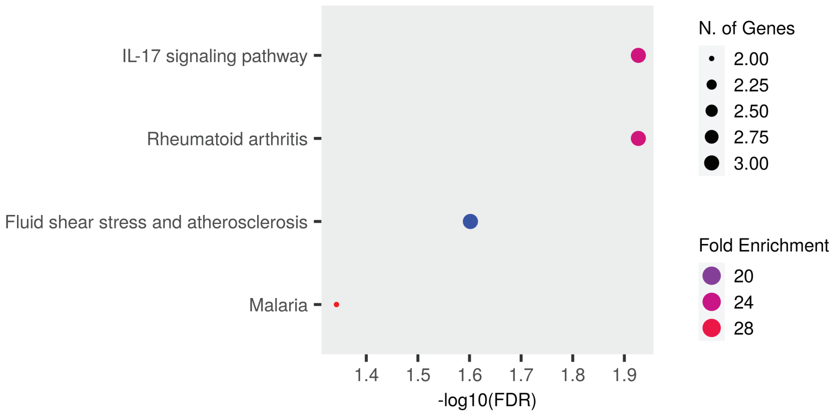


F Gene Type

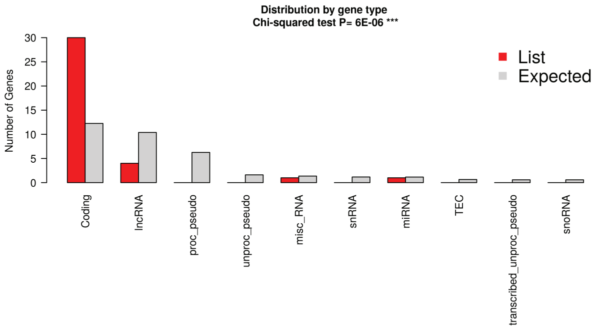
