## Extended figure 6 for "Brain Extracellular Matrix implications in multiple neurological disorders are revealed through a meta-analysis of transcriptional changes"

A Bipolar disorder Top DEGs

| Total No. of biploar disorder's studies analyzed: 7 | | |
|---|---|---|
| Gene Symbol | Name | Percentage of bipolar disorder's studies reported these DEGs |
| RASGRP1 | RAS guanyl releasing protein 1 | 28.6% |
| SLC35F1 | solute carrier family 35 member F1 | 28.6% |
| GNG12 | G protein subunit gamma 12 | 28.6% |
| IL4R | interleukin 4 receptor | 28.6% |
| MLC1 | modulator of VRAC current 1 | 28.6% |
| ETNPPL | ethanolamine-phosphate phospho-lyase | 28.6% |
| CA12 | carbonic anhydrase 12 | 28.6% |
| SENP6 | SUMO specific peptidase 6 | 28.6% |
| PLP1 | proteolipid protein 1 | 28.6% |
| TRAF3IP2 | TRAF3 interacting protein 2 | 28.6% |
| TGIF1 | TGFB induced factor homeobox 1 | 28.6% |
| TNFRSF11B | TNF receptor superfamily member 11b | 28.6% |
| HTR7 | 5-hydroxytryptamine receptor 7 | 28.6% |

B Cellular Components

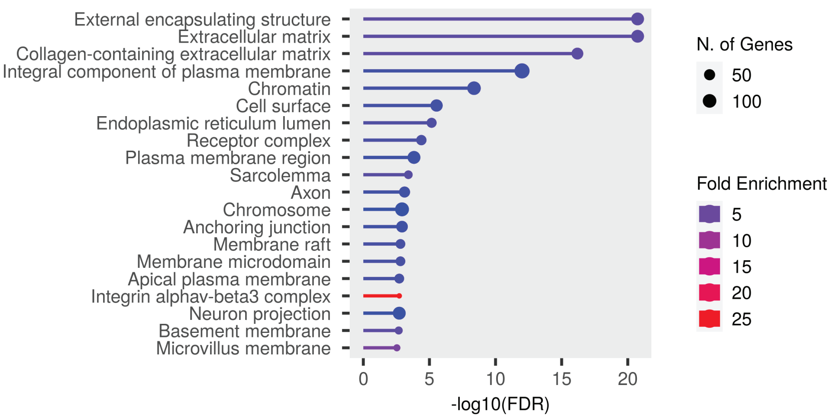


C Biological Processes

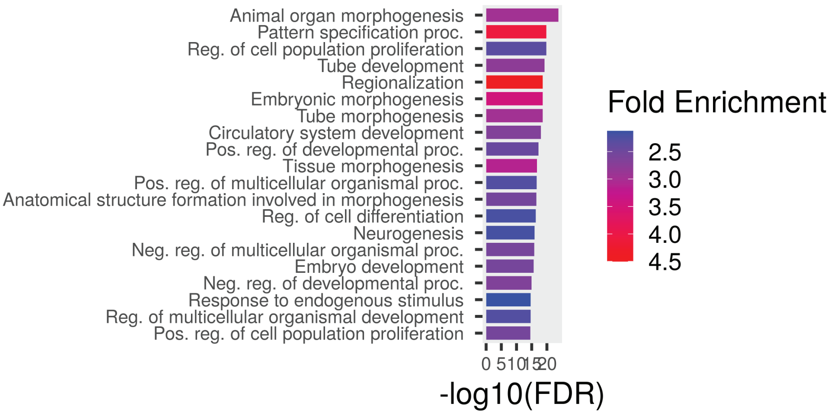


D Molecular Functions

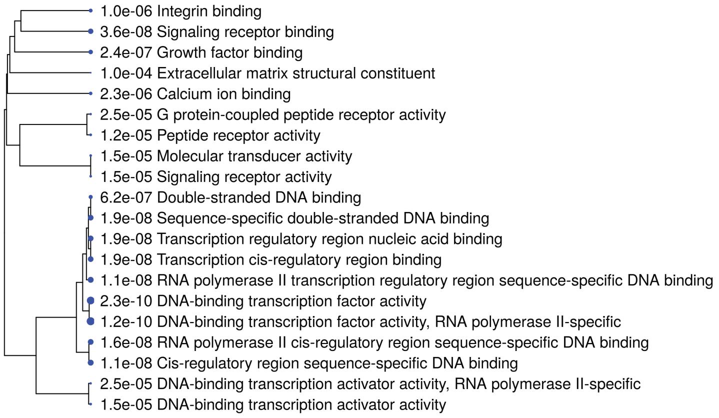


E KEGG

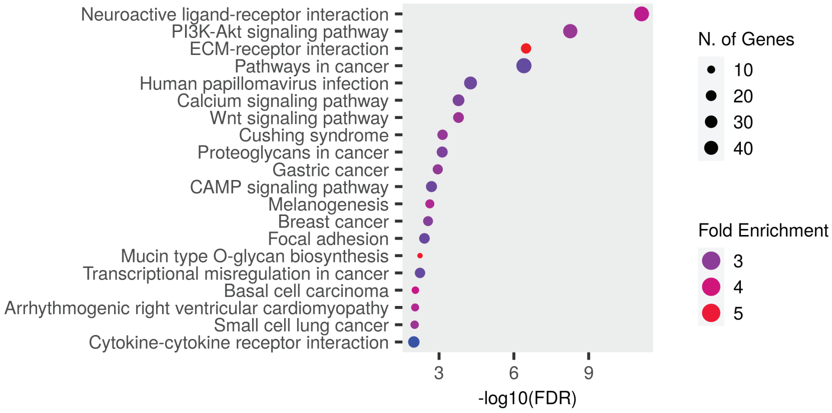


F Gene Type

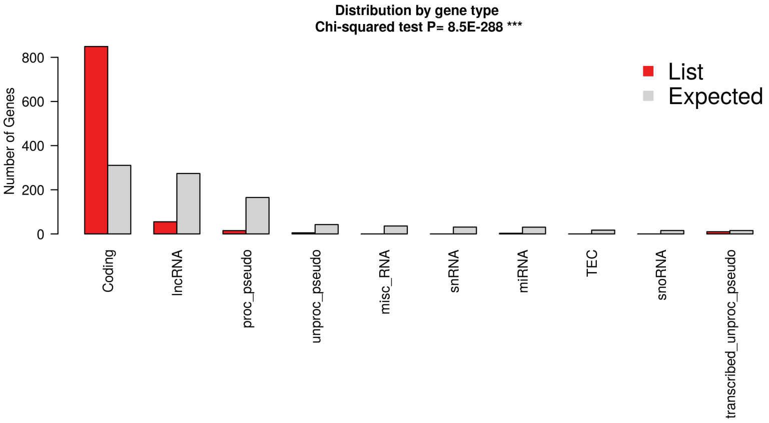
