## Supplementary figures and images for "Brain Extracellular Matrix implications in multiple neurological disorders are revealed through a meta-analysis of transcriptional changes"

### Extended figure 7

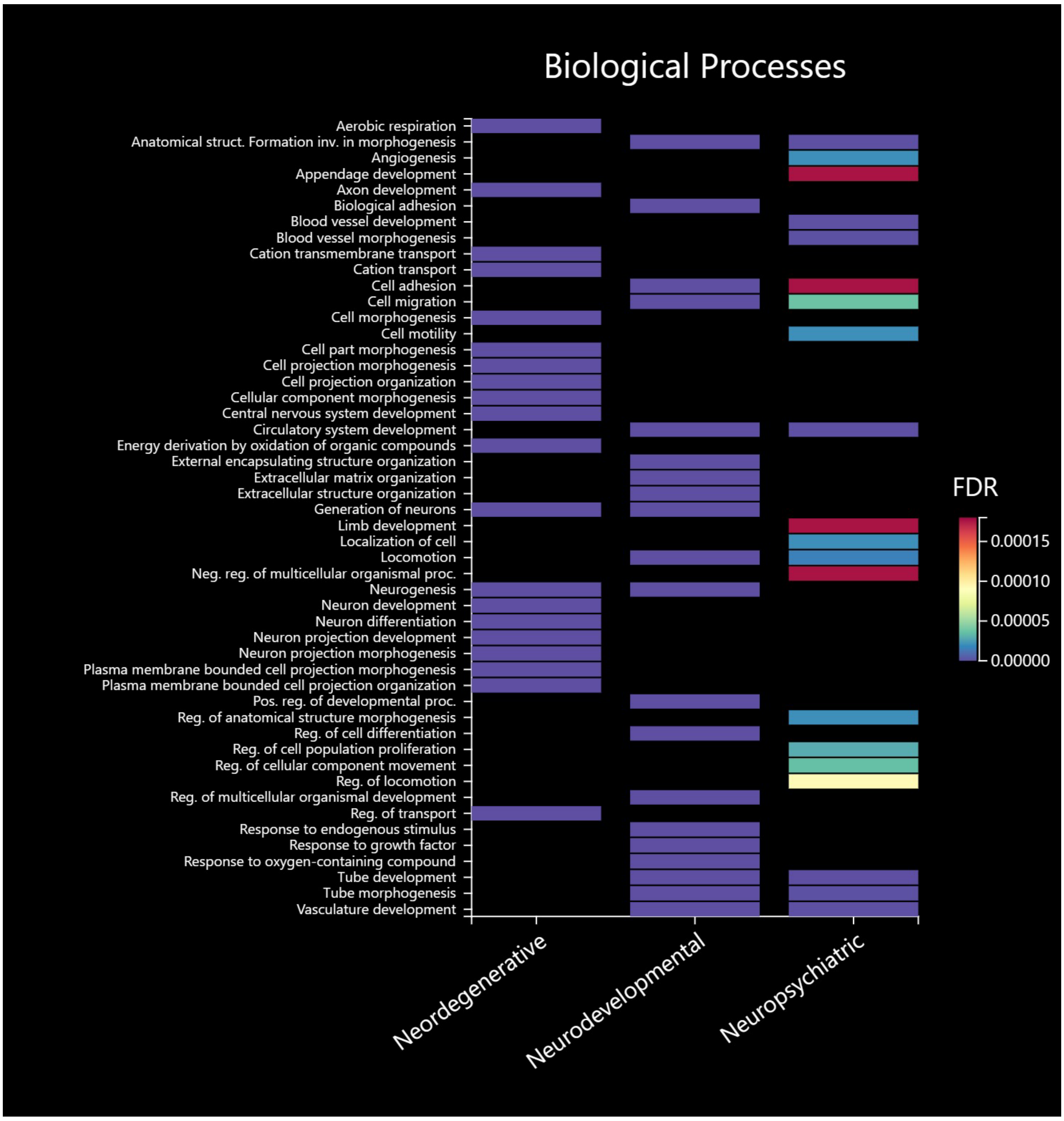
