## Supplementary S3 for "Brain Extracellular Matrix implications in multiple neurological disorders are revealed through a meta-analysis of transcriptional changes"

**Neurodegenerative disease analysis**

**Alzheimer's disease – dysregulation of Synaptic pathways along with AHNAK and SCG5 genes**

When comparing gene expression of postmortem brain tissues and neural organoids derived from Alzheimer's patients and healthy controls, we identified 8,382 significant DEGs (FDR < 0.05). Extended Figure 1A displays the top Alzheimer's disease reported DEGs, sorted by the percentage of overlapping genes. AHNAK and SCG5 genes had the highest ranking according to the percentage of overlapping genes among studies, with each appearing in 83.3% of the six relevant studies. Following them were 44 DEGs, each recurring in 66.7% of the six relevant studies, among them we found ATP5MC1, CSMD1, GAD1, HINT1, MDH1, NAP1L5, SNAP25, STMN2, and WWTR1 genes that were also highly reported in other neurodegenerative diseases examined in the current study.

Given the substantial number of dysregulated genes, Extended Figures 1B, C, D, and E display the GO analysis for 2,581 significant DEGs (FDR < 0.05) which were identified in more than one Alzheimer’s disease study out of the six relevant studies, all are coding genes (Extended Fig. 1F). Cellular components enrichment revealed several pathways including Synapse, Neuron projection, Axon, Somatodendritic compartment, Postsynapse, Cell body, Presynapse, and more (Extended Fig. 1B - cellular component pathways). When analyzing the enrichment for the biological processes, we found several pathways: Neurogenesis, Generation of neurons, Plasma membrane-bounded cell projection organization, Cell projection organization, Neuron differentiation, Trans-synaptic signaling, and Cellular component morphogenesis were among them (Extended Fig. 1C – biological process pathways). Molecular functions enrichment included Cytoskeletal protein binding, Cell adhesion molecule binding, Ion transmembrane transporter activity, Inorganic molecular entity transmembrane transporter activity, Inorganic cation transmembrane transporter activity, and Amyloid-beta binding, among others (Extended Fig. 1D - molecular function pathways). KEGG pathways were enriched for the following pathways: Pathways of neurodegeneration, Alzheimer's disease, Parkinson's disease, Huntington's disease, Prion disease, Amyotrophic lateral sclerosis, Focal adhesion, Regulation of actin cytoskeleton, and more (Extended Fig. 1E - KEGG).

**Parkinson's disease – dysregulated mitochondrial-related pathways**

When comparing gene expression of postmortem brain tissues and neurons derived from Parkinson’s patients and healthy controls, we identified 10,837 significant DEGs (FDR < 0.05). Extended figure 2A displays the top reported DEGs, sorted by the percentage of overlapping genes, including ACADVL, COL1A1, COL1A2, COL3A1, CPNE1, EEA1, FSTL1, GLIPR1, IFI16, KIAA1755, LSM4, NCAPD2, PAPPA, POU3F3, PTGDS, and TMEM107, each of them recurring in 80% of the five relevant studies.

Given the substantial number of dysregulated genes, Extended Figures 2B, C, D, and E display the GO analysis for 2,333 significant DEGs (FDR < 0.05) which were identified in more than one Parkinson’s disease study out of the five relevant studies, almost all are coding genes, and very few are of type lncRNA (Extended Fig. 2F).

Cellular components enrichment revealed several pathways including Neuron projection, Axon, Synapse, Inner mitochondrial membrane protein complex, Mitochondrial protein-containing complex, Organelle envelope, Envelope, Anchoring junction, Cell body, Mitochondrial membrane, Mitochondrial envelope, Mitochondrion, Mitochondrial respirasome, Focal adhesion, and more (Extended Fig. 2B - cellular component pathways). When analyzing the enrichment for the biological processes, we found several pathways: Neurogenesis, Generation of neurons, Neuron differentiation, cell projection organization, Plasma membrane-bounded Cell projection organization, and Response to endogenous stimulus were among them (Extended Fig. 2C – biological process pathways). The most significant Molecular functions enrichment included RNA binding, Cell adhesion molecule binding, Protein-containing complex binding, Cytoskeletal protein binding, and Electron transfer activity (Extended Fig. 2D - molecular function pathways). KEGG pathways were enriched for the following pathways: Parkinson's disease, Huntington's disease, Alzheimer's disease, Pathways of neurodegeneration, Prion disease, Amyotrophic lateral sclerosis, Oxidative phosphorylation, Diabetic cardiomyopathy, Metabolic pathways, PI3K-Akt signaling pathways, and more (Extended Fig. 2E - KEGG).

**Huntington's disease – dysregulation of urea metabolism**

When comparing gene expression of post-mortem brain tissues from Huntington’s patients and healthy controls, we identified 14,102 significant DEGs (FDR < 0.05). Extended figure 3A displays the most frequently reported DEGs, sorted by the percentage of overlapping genes, including BCL6, CEBPD, CRYM, FKBP5, GFAP, HTR2C, NEFM, PLOD2, and SLC14A1. Markedly, each of them was consistently identified in all 5 relevant studies related to Huntington’s disease. Protein-protein interaction (PPIs) network enrichment shows interactions between all 9 tops reported DEGs except the SLC14A1 gene (p <0.0023) (Extended Fig. 3B – Top genes PPIs networks). Surprisingly, SLC14A1, coding to solute carrier family 14 member 1 (Kidd blood group), is responsible for mediating the transport of urea. It has been reported that abnormal urea metabolism might serve as the initial biochemical disruption triggering neuropathogenesis in Huntington [1].

Given the substantial number of dysregulated genes, Extended Figures 3C, D, E, F, and G display the GO analysis for 2,898 significant DEGs (FDR < 0.05) which were identified in more than two Huntington’s disease studies out of the five relevant studies, all are coding genes (Extended Fig. 3C).

Cellular components enrichment revealed several pathways including Synapse, Neuron projection, Somatodendritic compartment, Postsynapse, Axon, Neuron to neuron synapse, Glutamatergic synapse, Dendritic tree, and more (Extended Fig. 3D - cellular component pathways). When analyzing the enrichment for the biological processes, we found several pathways: Neurogenesis, Neuron differentiation, Generation of neurons, Neuron projection development, Neuron development, Synaptic signaling, and cell morphogenesis were among them (Extended Fig. 3E – biological process pathways). The most significant Molecular functions enrichment included Cytoskeletal protein binding, Kinase binding, Calmodulin binding, Protein domain specific binding, and Actin binding (Extended Fig. 3F - molecular function pathways). KEGG pathways were enriched for the following pathways: Circadian entrainment, Glutamatergic synapse, CAMP signaling pathway, Oxytocin signaling pathway, Axon guidance, Long-term potentiation, MAPK signaling pathway, Pathways of neurodegeneration, Pathways in cancer, and more (Extended Fig. 3G – KEGG pathways).

**Neurodegenerative disease** **Discussion part**

**Alzheimer's disease**

Biological processes were significantly enriched with Neurogenesis and Generation of neuron pathways and Synaptic pathways were among the top dysregulated cellular components. The most frequently reported DEGs were SCG5, and AHNAK, each recurring in 83.3% of Alzheimer’s disease studies.

SCG5, coding for secretogranin V, also known as 7B2, is a secreted chaperone protein located in the extracellular region and nucleus, that prevents the aggregation of other secreted proteins, including proteins that are associated with neurodegenerative and metabolic disease. Chaplot et al. suggested that the potent chaperone action of SCG5 and its presence in a variety of extracellular protein deposits could have significance in neurodegenerative proteostatic mechanisms [2] [3] [4].

AHNAK is a large (700 kDa) structural scaffold protein, it has an essential role in developmental myelination processes, neuronal plasticity, and events related to neuroregeneration and neurodegeneration [3]. AHNAK was proposed as a potential biomarker of aging-related neurodegeneration [4].

**Huntington’s disease**

Biological processes were significantly enriched with Neurogenesis, Neuron differentiation, and Generation of neuron pathways. Synaptic pathways were among the top dysregulated cellular components. The most frequently reported DEGs included BCL6, CEBPD, CRYM, FKBP5, GFAP, HTR2C, NEFM, PLOD2, and SLC14A1. Markedly, each of them was consistently identified in all 5 relevant studies related to Huntington’s disease.

Significantly, Bailus et al. have conclusively shown that FKBP5 is involved in the progression of Huntington's disease. Inhibiting FKBP5 in human-derived neural stem cells (NSC) from individuals with Huntington's disease, as well as in diverse mouse models of HD, resulted in a decrease in mutant Huntingtin levels. Their findings propose a model wherein the reduction of FKBP5 levels or activity results in an increased autophagy process. This discovery positions FKBP5 as a potential therapeutic target	[5].

KEGG pathways were most significantly enriched with Circadian entrainment and Glutamatergic synapse pathways. Diago, E.B., et al. suggested that Huntington's disease patients may have a delayed sleep phase, that can be associated with their psychiatric symptoms [6]

**Parkinson’s disease**

Cellular components were significantly enriched with Synaptic pathways and mitochondrial-related pathways including Inner mitochondrial membrane protein complex, Mitochondrial protein-containing complex, Mitochondrial membrane, Mitochondrial envelope, Mitochondrion, Mitochondrial respirasome, and more. Biological process pathways were significantly enriched with Neurogenesis, Generation of neurons, and Neuron differentiation.

The top reported DEGs included ACADVL, COL1A1, COL1A2, COL3A1, CPNE1, EEA1, FSTL1, GLIPR1, IFI16, KIAA1755, LSM4, NCAPD2, PAPPA, POU3F3, PTGDS, and TMEM107 gene, each of them recurring in 80% of the five Parkinson’s disease studies.
