## Supplementary S4 for "Brain Extracellular Matrix implications in multiple neurological disorders are revealed through a meta-analysis of transcriptional changes"

**Neuropsychiatric disorders analysis**

**Schizophrenia – Dysregulation of Synaptic and ECM pathways**

When comparing gene expression of post-mortem brain tissues and neurons derived from Schizophrenia patients and healthy controls, we identified 3,955 significant DEGs (FDR < 0.05). Extended figure 4A displays the most frequently reported DEGs, sorted by the percentage of overlapping genes, including IFITM3, HSPA1A, and IFITM2, each of them recurring in 71.4% of the relevant studies, followed by IFITM1, HSPA1B, HSPB1, JMJD6, MEG3, and MKNK2, each recurring in 57.1% of the relevant studies.

Analysis of the cellular component enrichment reveals significant enrichment for the following pathways: Synapse, Neuron projection, Axon, Somatodendritic compartment, Extracellular matrix, External encapsulating structure, Collagen-containing extracellular matrix, and more (Extended Fig. 4B - cellular component pathways). When analyzing the enrichment for the biological processes we found that the most significant pathways were Circulatory system development, Neurogenesis, Locomotion, Regulation of anatomical structure morphogenesis, Generation of neurons, Cell projection organization, and Neuron projection development (Extended Fig. 4C – biological process pathways). Molecular Functions were mostly enriched for Cell adhesion molecule binding, Signaling receptor binding, Calcium ion binding, and Ion transmembrane transporter activity (Fig. 4D – molecular functions). KEGG pathways were enriched for the following pathways: Pathways of neurodegeneration, Amyotrophic lateral sclerosis, Oxidative phosphorylation, MAPK signaling pathways, Focal adhesion, PI3K-Akt signaling pathways, Metabolic pathways, and more (Extended Fig. 4E – KEGG pathways). Almost all the dysregulated genes were coding genes, and only a few were of type incRNA (Extended Fig. 4F – Gene type).

**Major Depression**

When comparing gene expression of post-mortem brain tissues from major depression patients and healthy controls, we identified 12,221 significant DEGs (FDR < 0.05). Extended Figure 5A displays the most frequently reported DEGs, sorted by the percentage of overlapping genes. FOS gene had the highest ranking according to the percentage of reporting, recurring in 75% of Major depression studies, followed by additional 42 DEGs recurring in 50% of the relevant studies, including HLA-A, TM4SF1, IFI6, RGS1, FAM118A, and HSPA1A genes that were also highly reported in other neurological disorders examined in the current study.

Given the substantial number of dysregulated genes, Extended Figures 5, B, C, D, E, and F, display the GO analysis for the 43 significant DEGs (FDR < 0.05) which were identified in more than one major depression study out of four relevant studies (very low recurrence compared to a total of 12,221 DEGs identified for major depression)

Cellular components enrichment didn't show significant pathways, but when grouping the 43 DEGs by functional cellular component categories defined by high-level GO terms, we observe that 14 of the 43 DEGs belong to the Extracellular region, 11 to the Extracellular space, 6 to the Organelle membrane, 6 to the Extracellular organelle, 5 to the Chromatin organelle, and 4 to Cell projection, with some overlapping between categories (Extended Fig. 5B). When analyzing the enrichment for the biological processes, we found several pathways: [Positive regulation of cell death](http://amigo.geneontology.org/amigo/term/GO:0010942), [Skeletal muscle cell differentiation](http://amigo.geneontology.org/amigo/term/GO:0035914), [progesterone biosynthetic process](http://amigo.geneontology.org/amigo/term/GO:2000182), [Cytokine-mediated signaling pathway](http://amigo.geneontology.org/amigo/term/GO:0019221), [Response to cytokine](http://amigo.geneontology.org/amigo/term/GO:0034097), Cellular response to cytokines stimulus, Response to external biotic stimulus, and more (Extended Fig. 5C – biological process pathways). Molecular functions enrichment included [DNA-binding transcription activator activity](http://amigo.geneontology.org/amigo/term/GO:0001216) RNA polymerase ||-specific, and [DNA-binding transcription activator activity](http://amigo.geneontology.org/amigo/term/GO:0001216) (Extended Fig. 5D - molecular function pathways). KEGG pathways were enriched for the following pathways: [IL-17 signaling pathway](http://www.genome.jp/kegg-bin/show_pathway?hsa04657) (The interleukin 17 family, a subset of cytokines consisting of IL-17A-F, plays crucial roles in both acute and chronic inflammatory responses), [Rheumatoid arthritis](http://www.genome.jp/kegg-bin/show_pathway?hsa05323) (a chronic autoimmune joint disease where persistent inflammation affects bone remodeling leading to progressive bone destruction), [Fluid shear stress and atherosclerosis](http://www.genome.jp/kegg-bin/show_pathway?hsa05418), and [Malaria](http://www.genome.jp/kegg-bin/show_pathway?hsa05144) (Extended Fig. 5E – KEGG pathways). 30 of the 42 dysregulated genes were coding genes, the rest were of lncRNA, miRNA, and misc RNA.

**Bipolar disorder**

When comparing gene expression of post-mortem brain tissues and neurons derived from bipolar disorder patients and healthy controls, we identified 993 significant DEGs (FDR < 0.05), the lowest number of DEGs among all the 7 disorders examined in the current study. Extended figure 6A displays the most frequently reported DEGs, sorted by the percentage of overlapping genes, including CA12, ETNPPL, GNG12, HTR7, IL4R, MLC1, PLP1, RASGRP1, SENP6, SLC35F1, TGIF1, TNFRSF11B, AND TRAF3IP2 each recurring in 28.6% of the relevant studies, significantly lower reporting rate than all other disorders, with maximum recurrence of 2 out of 7 bipolar disorder studies.

Analysis of the cellular component enrichment reveals that the 993 DEGs showed significant enrichment for the following pathways: Extracellular Matrix, External encapsulating structure, Collagen-containing extracellular matrix, Integral component of the plasma membrane, and more (Extended Fig. 6B - cellular components pathways). When analyzing the enrichment for the biological processes we found that the most significant pathways were Animal organ morphogenesis, Pattern specification process, [Regulation of cell population proliferation](http://amigo.geneontology.org/amigo/term/GO:0042127), and [Tube development](http://amigo.geneontology.org/amigo/term/GO:0035295) (Extended Fig. 6C – biological processes). Molecular Functions were enriched for [DNA-binding transcription factor activity - RNA polymerase II-specific](http://amigo.geneontology.org/amigo/term/GO:0000981), and DNA-binding transcription factor activity, [RNA polymerase II transcription regulatory region sequence-specific DNA binding](http://amigo.geneontology.org/amigo/term/GO:0000977), [Cis-regulatory region sequence-specific DNA binding](http://amigo.geneontology.org/amigo/term/GO:0000987), [RNA polymerase II cis-regulatory region sequence-specific DNA binding](http://amigo.geneontology.org/amigo/term/GO:0000978), and more (Extended Fig. 6D - molecular functions). KEGG pathways were enriched for the following pathways: [Neuroactive ligand-receptor interaction](http://www.genome.jp/kegg-bin/show_pathway?hsa04080), [PI3K-Akt signaling pathway](http://www.genome.jp/kegg-bin/show_pathway?hsa04151), [ECM-receptor interaction](http://www.genome.jp/kegg-bin/show_pathway?hsa04512), and more (Extended Fig. 6E). The characteristics of the dysregulated genes are displayed in Extended Fig 4F, almost all the dysregulated genes were coding genes, less than 10% were of type lncRNA, and very few were of type proc_pseudo, transcribed_unproc_pseudo, and unproc_pseudo.

**Neuropsychiatric Disorders Discussion part**

**Bipolar-disorder**

We identified 13 significant DEGs (FDR<0.05) that appeared in more than one bipolar disorder study out of the four bipolar disorder studies examined in the current study.

Cellular components were significantly enriched with ECM pathways. KEGG pathways were significantly enriched for [Neuroactive ligand-receptor interaction](http://www.genome.jp/kegg-bin/show_pathway?hsa04080), [PI3K-Akt signaling pathway](http://www.genome.jp/kegg-bin/show_pathway?hsa04151), [ECM-receptor interaction](http://www.genome.jp/kegg-bin/show_pathway?hsa04512), and more. Molecular functions were highly enriched with [DNA-binding transcription factor activity - RNA polymerase II-specific](http://amigo.geneontology.org/amigo/term/GO:0000981), and DNA-binding transcription factor activity, [RNA polymerase II transcription regulatory region sequence-specific DNA binding](http://amigo.geneontology.org/amigo/term/GO:0000977), [Cis-regulatory region sequence-specific DNA binding](http://amigo.geneontology.org/amigo/term/GO:0000987), [RNA polymerase II cis-regulatory region sequence-specific DNA binding](http://amigo.geneontology.org/amigo/term/GO:0000978), and more. Interestingly, bipolar disorder demonstrated the lowest number of significant DEGs identified among all the 7 disorders examined in the current study and also showed the lowest percentage of gene recurrence. Specifically, out of 993 genes reported to be differentially expressed between bipolar disorder patients and a healthy control group, only 13 genes were found in more than 1 study, and none of them appeared in more than 2 out of the 7 bipolar disorder studies analyzed in the current study. These 13 genes that exhibit recurrence were: RASGRP1, SLC35F1, GNG12, IL4R, MLC1, ETNPPL, CA12, SENP6, PLP1, TRAF3IP2, TGIF1, TNFRSF11B, and HTR7.

One possible explanation for the low recurrence could be the substantial variability observed among bipolar patients, encompassing distinct cellular parameters between lithium-responsive patients-derived neurons and non-responsive patients-derived neurons, leading to the emergence of discrete neuronal subgroups that predict patients' responsiveness to lithium [1]. Additionally, distinct epigenetic mechanisms influencing gene expression categorize individuals with bipolar disorder into two groups: those with suicidal tendencies and those with non-suicidal tendencies [2].

**Major Depression**

biological processes pathways were significantly enriched with [Positive regulation of cell death](http://amigo.geneontology.org/amigo/term/GO:0010942), [Skeletal muscle cell differentiation](http://amigo.geneontology.org/amigo/term/GO:0035914), [progesterone biosynthetic process](http://amigo.geneontology.org/amigo/term/GO:2000182), [Cytokine-mediated signaling pathway](http://amigo.geneontology.org/amigo/term/GO:0019221), [Response to cytokine](http://amigo.geneontology.org/amigo/term/GO:0034097), Cellular response to cytokines stimulus, Response to external biotic stimulus, and more (Extended Fig. 5C – biological process pathways). Molecular functions enrichment included [DNA-binding transcription activator activity](http://amigo.geneontology.org/amigo/term/GO:0001216) RNA polymerase ||-specific, and [DNA-binding transcription activator activity](http://amigo.geneontology.org/amigo/term/GO:0001216) (Extended Fig. 5D - molecular function pathways). KEGG pathways were enriched for the following pathways: [IL-17 signaling pathway](http://www.genome.jp/kegg-bin/show_pathway?hsa04657) (The interleukin 17 family, a subset of cytokines consisting of IL-17A-F, plays crucial roles in both acute and chronic inflammatory responses), [Rheumatoid arthritis](http://www.genome.jp/kegg-bin/show_pathway?hsa05323) (a chronic autoimmune joint disease where persistent inflammation affects bone remodeling leading to progressive bone destruction), [Fluid shear stress and atherosclerosis](http://www.genome.jp/kegg-bin/show_pathway?hsa05418), and [Malaria](http://www.genome.jp/kegg-bin/show_pathway?hsa05144)

FOS gene had the highest ranking according to the percentage of reporting, recurring in 75% of Major depression studies, followed by additional 42 DEGs recurring in 50% of the relevant studies, including HLA-A, TM4SF1, IFI6, RGS1, FAM118A, and HSPA1A genes that were also highly reported in other neurological disorders examined in the current study.

**Schizophrenia**

we identified 3,955 significant DEGs (FDR < 0.05). The most frequently reported DEGs, included HSPA1A, IFITM2, and IFITM3, each of them recurring in 71.4% of the relevant studies, followed by IFITM1, HSPA1B, HSPB1, JMJD6, MEG3, and MKNK2, each recurring in 57.1% of the relevant studies.

Biological processes were significantly enriched with Circulatory system development, Neurogenesis, Locomotion, and more.

HSPB1, encoding for heat shock protein family B (small) member 1, also known as HSP27, contributes significantly to cell survival in stressful conditions, and plays an important role during the CNS development, aligning with the neurodevelopmental hypothesis of schizophrenia, implicates abnormal or disrupted neural growth during embryonic development [3]. HSPB1 is upregulated in neuronal progenitors undergoing differentiation [4]. Abnormal expression of HSPB1 was previously demonstrated in schizophrenia patients [5]. In addition, HSPB1 can interact with a broad range of substrates. This interaction prevents the creation and elongation of amyloid fibrils, inhibits the aggregation of misfolded polypeptides, and indirectly facilitates their refolding or degradation [6], and has been noted to associate with protein aggregates in various neurodegenerative disorders [5].

Notably, HSPB1 was also highly reported in neurodevelopmental studies, recurring in 57.1% of relevant studies, the same as schizophrenia’s reporting rate.
